# Control of alveolar macrophage differentiation by Siah1a/2 ubiquitin ligases limits carcinogen-induced lung adenocarcinoma

**DOI:** 10.1101/2022.09.14.508032

**Authors:** Marzia Scortegagna, Yuanning Du, Linda M. Bradley, Kun Wang, Eytan Ruppin, Rabi Murad, Ze’ev A. Ronai

## Abstract

Tumor microenvironment components, including T and myeloid cells, play important roles in lung adenocarcinoma (LADC) progression and response to therapy. Here, we identify a role for Siah1a/2 ubiquitin ligases in controlling alveolar macrophage (AM) differentiation and urethane-induced LADC. Genetic ablation of Siah1a/2 in AMs enriched their immature state, coinciding with increased pro-tumorigenic and inflammatory gene signatures and abundance of Siah1a/2 substrates NRF2 and β-catenin. Urethane administration to mice enriched the population of monocytic and immature-like AMs, which were more prevalent in the lungs of mice carrying Siah1a/2-ablated macrophages. While resembling transitional profibrotic macrophages often seen in lung fibrosis, enrichment of immature AMs coincided with the development of more frequent and larger lung tumors in urethane-treated mice harboring Siah1a/2 ablated macrophages compared with controls. Gene expression signature of Siah1a/2 ablated immature-like AMs is associated with increased infiltration of CD14+ immune cells and worse survival of LADC patients. Our findings identify Siah1a/2 as gatekeepers of cancer development by controlling AM differentiation and profibrotic phenotypes contributing to carcinogen-induced lung cancer.

## Introduction

Our understanding of how the immune system controls tumor development, progression, and therapy response has prompted development of therapies to modify and exploit select immune system components ^1, 2^. Complex immune system/tumor cross-talk is in part mediated by tumor microenvironment (TME) components, now known to be important determinants of immune cell activation and infiltration. Specific immune cell components, including cytotoxic and regulatory T cells and dendritic cells, function in the TME and are implicated in T cell infiltration and activation and therefore in control of tumor growth ^3-4^. Other TME components include fibroblasts and endothelial cells ^5,6^ as well as macrophages, which are less studied myeloid cells recently recognized as controlling tumor growth ^7,8^. Here we focus on the role of macrophages in lung cancer development using a unique genetic model in which the ubiquitin ligases Siah1a/2 are ablated in macrophages.

Approximately 85% of lung cancers are non-small cell lung cancer (NSCLC), with lung adenocarcinoma (LADC) and squamous cell carcinoma the most common subtypes ^9^. In response to chemical carcinogens (i.e., urethane), mice develop pulmonary adenocarcinoma resembling human adenocarcinoma harboring KRAS mutations ^10^. Several studies have reported the importance of the lung microenvironment, including immune system components, in lung cancer development, and many highlight the abundance of macrophages and their possible contribution to its development ^11,12^. Two macrophage populations, alveolar macrophages (AMs) and interstitial macrophages (IMs), are abundant in normal lung: AMs are in close contact with type I and II epithelial cells in alveoli, while IMs are found in parenchyma between the vascular endothelium and the alveolar epithelium ^13,14^. AMs are derived from fetal monocytes that populate alveoli soon after birth and self-renew independently of bone marrow contribution ^15,16^. Fetal monocytes accumulate in the developing mouse lung at E14.5 and differentiate into immature (F4/80^+^, CD11c^+^, CD11b intermediate, and SiglecF low) AMs ^17,18^. They then mature postnatally into AMs (CD11c^+^ F4/80^+^ SiglecF^+^ and CD11b low), a process requiring GM-CSF-PPRγ signaling ^16^. After inflammatory insult, bone marrow-derived cells can differentiate into AMs ^16^. Notably, microenvironmental factors, including oxygen tension, glucose supply, and exposure to surfactant-rich fluid, all of which fluctuate during acute infection or chronic inflammation, as well as communication with alveolar epithelial cells, define AM development and function ^19,20^. Relevant to cancer, previous studies identified correlations between density of tumor-associated macrophages (TAMs), their abundance in stroma, and M2 polarization with worse patient survival ^12,11^. Of note, AMs obtained by bronchoalveolar lavage from patients with lung cancer exhibit reduced phagocytic capabilities ^21^, although their role in lung cancer progression is still debated. This study extends our earlier analysis of Siah2 in anti-tumor immunity. *Siah2* mutant mice show decreased melanoma tumor growth, an outcome most notable in the context of immunogenic melanomas. Mechanistically, Siah2 reportedly regulates activity of T cells, primarily Tregs, by controlling their proliferation and infiltration of melanoma tumors ^22^. Likewise, treatment of *Siah2*^*KO*^ mice with anti-PD1 therapy effectively blocked development of therapy-resistant (“cold”) melanoma ^22^. These observations established the importance of Siah2 in the TME and extended earlier studies that established Siah2 importance in tumor-intrinsic functions. Previously, in analyzing melanoma infiltration by immune cells we developed Siah2 KO mice and identified enrichment of macrophage sub-populations. Here we analyzed Siah1a/2 function in macrophages by conditionally ablating Siah1a/2 genes in macrophage cells. Mice deficient in Siah1a/2 in macrophages harbored AMs in an immature-like state, and those cells exhibited a unique inflammatory signature, which was enhanced in mice treated with the carcinogen urethane. Urethane-treated mice also exhibited more frequent and larger lung tumors than WT controls, suggesting that Siah1a/2 serve as gatekeepers of macrophage ability to control lung cancer development.

## Results

### Siah1a and Siah2 are required for AM terminal differentiation and maturation

To establish mice with conditional loss of Siah1a and Siah2 (Siah1a/2) in myeloid cells, we generated *Siah1a*^*f/f*^*::Siah2*^*f/f*^ mice and then crossed them with macrophage-specific Cre (*Lyz2*^*Cre*^) mice, to ablate Siah1a/2 in macrophages (*cSiah1a/2*^*f/f*^*::Lyz2*^*Cre*^). We then characterized macrophages in spleen, heart, and kidney of *cSiah1a/2*^*f/f*^*::Lyz2*^*Cre*^ and WT mice focusing on CD45^+^ cells enriched for MΘ markers (CD11b^+^ and F480^+^). We observed that the frequency of these cells in each organ was comparable in both genotypes (Sup. Figure 1A). We next used markers of MΘ subtypes to FACS-sort and characterize AM and IM populations in mouse lung tissues. The overall % of either CD45^+^ cells or of IMs (CD11b^+^ F4/80^+^) was comparable in lungs of both genotypes (Figure 1A and Sup. Figure 1B). However, Siah1a/2 loss in macrophages decreased AM (CD11b low F480^+^) frequency in lung relative to WT cells (Figure 1B, F). This decrease was further reflected in the fraction of mature AMs (CD11b low F4/80^+^ CD11c^+^ SiglecF^+^) (Figures 1C, 1F). These findings suggest that AMs are the primary MΘ population whose differentiation requires Siah1a/2. Of note, most CD11b low F480^+^ cells were positive for the mannose receptor CD206^+^ in both WT and *cSiah1a/2*^*f/f*^*::Lyz2*^*Cre*^ genotypes (Sup. Figure 1C), confirming the high expression of CD206 in AMs.

**Figure 1.**
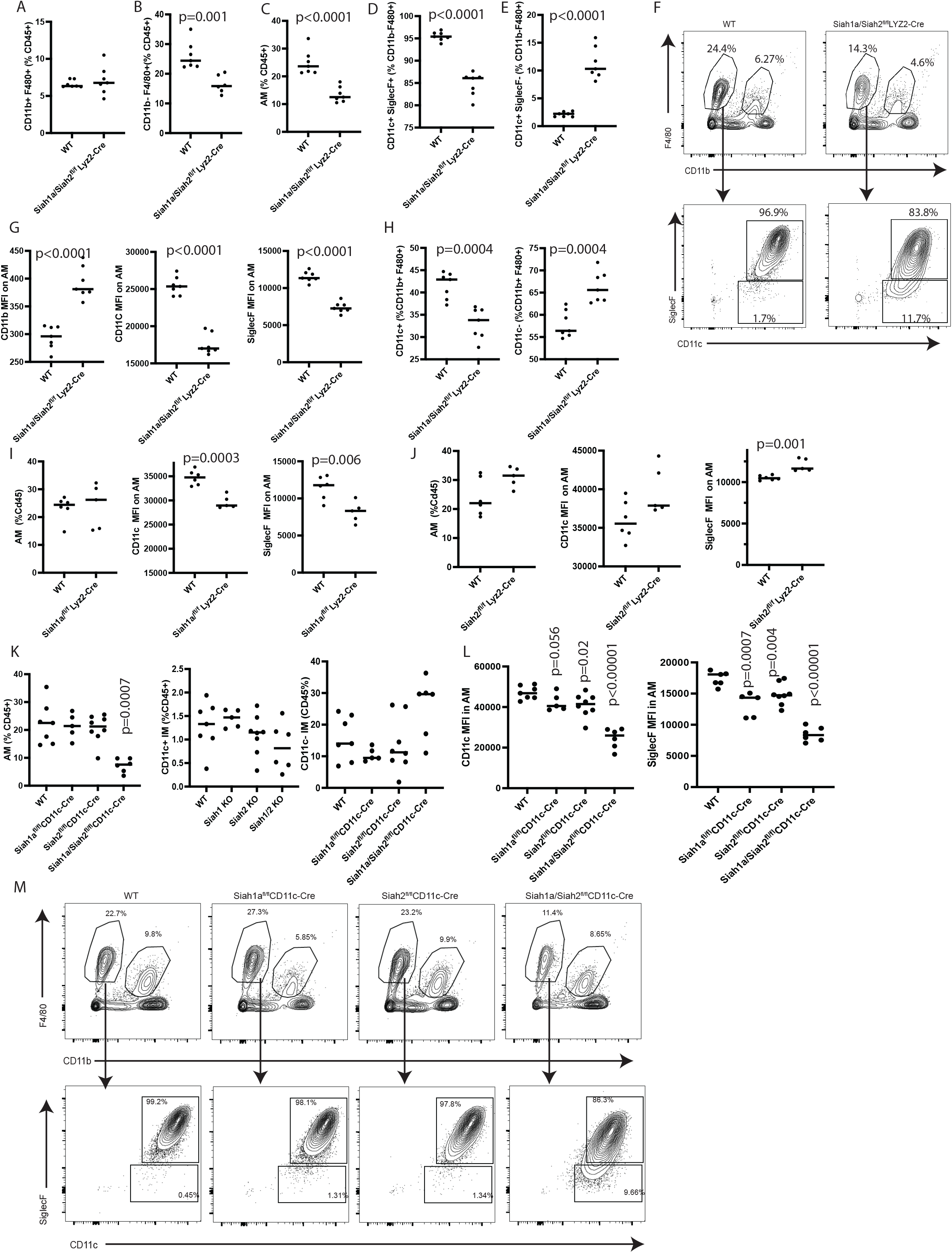
Siah1a and Siah2 are required for AM terminal differentiation and maturation. **A-C** Quantification of CD11b^+^ F480^+^ (A) and CD11b low F480^+^ (B) immune cells, and AMs (CD11b low F480^+^ CD11c^+^ SiglecF^+^) (C) from lungs of *WT* and *cSiah1a/2*^*f/f*^*::Lyz2*^*Cre*^ mice at homeostatic state. Quantification is shown as a percentage of indicated immune cells among CD45.2^+^ cells. n=7 for both genotypes. **D, E** Quantification of AMs positive for SiglecF and CD11c (D) or CD11c^+^ and SiglecF low (E) from lungs of *WT* and *cSiah1a/2*^*f/f*^*::Lyz2*^*Cre*^ mice. Quantification is shown as a frequency of immune cells among CD11b low F/480^+^ cells. n=7 for both genotypes. **F** FACS analysis of expression of CD11c^+^ SiglecF^+^ cells by a gated subpopulation (CD11b low F480^+^ cells) of CD45.2^+^ cells from *WT* and *cSiah1a/2*^*f/f*^*::Lyz2*^*Cre*^ lungs. **G** Expression of CD11b, CD11c and SiglecF on AMs (CD11b low F480+ CD11c^+^ SiglecF^+^ cells) based on flow cytometry analysis of lungs from *WT* and *cSiah1a/2*^*f/f*^*::Lyz2*^*Cre*^ mice. n=7 for both genotypes. **H** CD11c+ and CD11c-cell frequency within the population of *WT* and *cSiah1a/2*^*f/f*^*::Lyz2*^*Cre*^ IMs (CD11b+ F4/80+). n=7 for both genotypes. **I, J** Quantification of AMs and CD11c^+^ and CD11c^-^ IMs from lungs of *WT* and *cSiah1a*^*f/f*^*::Lyz2*^*Cre*^ (I) and *cSiah2*^*f/f*^*::Lyz2*^*Cre*^ (J) mice at a homeostatic state. Shown is a percentage of indicated immune cells among CD45.2^+^ cells. n=6 for WT; n=5 for *cSiah1a*^*f/f*^*::Lyz2*^*Cre*^; n=6 for WT; n=5 for *cSiah2*^*f/f*^*::Lyz2*^*Cre*^. **K** Quantification of AMs and CD11c+ and CD11c-IMs from *WT* and *cSiah1a*^*f/f*^*::CD11c*^*Cre*^, *cSiah1a/2*^*f/f*^*::CD11c*^*Cre*^ and *cSiah2*^*f/f*^*::CD11c*^*Cre*^ mice. Quantification is shown as a percentage of indicated immune cells among CD45.2^+^ cells. n=7 for WT; n=5 for *cSiah1a*^*f/f*^*::CD11c*^*Cre*^; n=8 for *cSiah2*^*f/f*^*::CD11c*^*Cre*^; n=6 for *cSiah1a/2*^*f/f*^*::CD11c*^*Cre*^. **L** CD11c, CD206, and SiglecF expression on AMs (CD11b low F480^+^ CD11c^+^ SiglecF+ cells), based on flow cytometry analysis of lungs from *WT* and *cSiah1a*^*f/f*^*::CD11c*^*Cre*^, *cSiah2*^*f/f*^*::CD11c*^*Cre*^ and *cSiah1a/2*^*f/f*^*::CD11c*^*Cre*^ mice. n=7 for WT; n=5 for *cSiah1a*^*f/f*^*::CD11c*^*Cre*^; n=8 for *cSiah2*^*f/f*^*::CD11c*^*Cre*^; n=6 for *cSiah1a/2*^*f/f*^*::CD11c*^*Cre*^. **M** Representative flow cytometry showing expression of CD11c^+^ SiglecF^+^ cells by a gated subpopulation (CD11b low F480^+^ cells) of CD45.2+ cells from *WT* and *cSiah1a*^*f/f*^*::CD11c*^*Cre*^, *cSiah2*^*f/f*^*::CD11c*^*Cre*^ and *cSiah1a/2*^*f/f*^*::CD11c*^*Cre*^ lungs. Data were analyzed by unpaired t-test.

Accordingly, in WT mice, most CD11b low F480^+^ cells were positive for other markers of mature AMs (SiglecF and CD11c); however, in *cSiah1a/2*^*f/f*^*::Lyz2*^*Cre*^ mice a smaller fraction was positive for mature macrophage markers (Figure 1D). Correspondingly, the frequency of immature AMs (CD11b low F480^+^CD11c^+^ SiglecF^-^) in the AM population was 4 times higher in *cSiah1a/2*^*f/f*^*::Lyz2*^*Cre*^ relative to WT mice (Figure 1E).

Notably, in the mature (CD11b low F4/80^+^ CD11c^+^ SiglecF^+^) AM population from *cSiah1a/2*^*f/f*^*::Lyz2*^*Cre*^ mice, we observed a significant decrease in CD11c- and SiglecF-expressing cells and an increase in CD11b expression relative to that seen in WT mice (Figure 1G), confirming that Siah1a/2-deficient AMs exhibit a relatively immature-like phenotype compared to WT mice. Nonetheless, expression of F4/80, MHCI, CD80 and CD206 in AMs was comparable in macrophages from both genotypes (Sup. Figure 1D), suggesting that changes seen in CD11c and SiglecF expression in AMs reflect their differentiation status.

Decreased CD11c expression seen in Siah1a/2-deficient AMs coupled with knowledge that a subpopulation of IMs express CD11c pointed to the possibility that Siah1a/2 affects CD11c^+^ IMs. Indeed, we observed that the frequency of CD11c^+^ IMs slightly but significantly decreased (43% of CD45^+^ cells in WT compared to 33% in Siah1a/2-deleted cells), while the frequency of CD11c^-^ IMs significantly increased among IMs from *cSiah1a/2*^*f/f*^*::Lyz2*^*Cre*^ compared to WT mice (Figure 1H). These changes did not alter CD206 frequency, and the expression of CD11c within the CD11c^+^ IM population was comparable in WT and *cSiah1a/2*^*f/f*^*::Lyz2*^*Cre*^ cells (Sup. Figure 1E). Importantly, we observed no changes in frequency of other immune cell populations, including neutrophils, CD4, CD8, or B cells, or monocytes, except for a significant decrease in NK1.1^+^ cells (Sup Figure 1F). Overall, these data suggest that Siah1a and Siah2 are required for AM cell differentiation to a fully mature state.

To assess the relative contribution of each ligase to phenotypes described above, we generated mice in which a single Siah gene was ablated in macrophages *(Siah1a*^*f/f*^*:Lyz2*^*Cre*^ and *Siah2*^*f/f*^:*Lyz2*^*Cre*^) and then subjected lungs from these mice to FACS analysis. AM frequency among CD45^+^ cells was comparable in mice harboring WT, Siah1- or Siah2-deficient macrophages, although we observed a slight but significant decrease of CD11c and SiglecF expression on AMs from *cSiah1a*^*f/f*^*::Lyz2*^*Cre*^ relative to WT mice and a slight but significant increase of SiglecF expression on AMs from *cSiah2*^*f/f*^*::Lyz2*^*Cre*^ relative to WT mice (Figure 1I and J). Notably, these differences were less pronounced than those seen in DKO (*cSiah1a/2*^*f/f*^*::Lyz2*^*Cre*^) macrophages, suggesting that Siah2 can augment changes triggered by Siah1a in AM. While the CD11b^+^ F480^+^ population was comparable among WT and Siah1a-depleted cells, we observed a slight but significant decrease of this population in Siah2 depleted cells, when compared to WT. We also observed a slight but significant decrease in CD11c^+^ IMs, along with a similar decrease in CD11c^-^ IMs among *cSiah1a*^*f/f*^*::Lyz2*^*Cre*^ as opposed to WT genotypes. (Sup figure 1 G). These data indicate that both Siah1a and Siah2 are necessary for AM maturation.

*Lyz2 C*re-induced gene deletion can occur embryonically ^16^. To determine whether phenotypes described here emerge pre- or post-natal, we performed comparable analysis in a mouse model in which Siah1/2 ablation is catalyzed by CD11c-Cre, a driver expressed in AMs at its highest levels after birth ^16^. This approach allowed us to confirm Siah1a/2 function in maintenance of AM homeostasis and potentially exclude a cell-intrinsic role of Siah1a/2 in neutrophils (which also express Lyz2). Indeed, crossing of CD11c^Cre^ with *Siah1a*^*f/f*^ *:Siah2*^*f/f*^ mice ablated Siah1a and Siah2 in AMs, CD11c^+^ IMs and dendritic cells. As we observed in *Lyz*^*Cre*^ mice, CD45^+^ cell frequency in lung was comparable in WT and mutant mice (Sup Figure 1H). Likewise, Siah1a/Siah2 ablation via CD11C^Cre^ significantly decreased the frequency of AMs and CD11c^+^ IMs, while increased the frequency of CD11c^-^ IMs (Figure 1K, M). CD11c, SiglecF and CD206 expression also significantly decreased in AMs from *cSiah1a/2*^*f/f*^*::CD11c*^*Cre*^ compared to WT lungs (Figure 1L), phenotypes comparable to those seen in analysis using the Lyz2 driver. Of interest, single deletion of Siah1a or Siah2 in CD11c^+^ cells did not alter frequency of AMs, CD11c^+^ IMs or CD11c^-^ IMs (figure 1K, M). Within the AM population, CD11c and CD206 expression was comparable in WT and Siah1a- or Siah2-deleted cells, while we observed a small but significant decrease in SiglecF expression in Siah1a-deleted relative to WT cells (Figure 1L). Here, by using a second mouse model, we confirmed the role of Siah1a/2 in AM maturation and differentiation.

Dendritic cell frequency or that of other immune cell populations (including CD4 or CD8 cells, neutrophils, or B cells) was comparable in WT and Siah1a or Siah2 singly- or doubly-deleted cells (Supplemental Figure I). We also confirmed a significant decrease in frequency of NK1.1-positive cells seen in both *cSiah1a/2*^*f/f*^*::Lyz2*^*Cre*^ and in *cSiah1a/2*^*f/f*^*::CD11c*^*Cre*^ CD45^+^ cells of the lungs (Sup. Figure 1J), suggesting that Siah1a and Siah2 expression in the myeloid population is responsible for NK1.1 cell recruitment.

### Siah1a and Siah2 regulates AM maturation of fetal monocytes to AMs

To determine the possible role of Siah1a/2 in development of fetal to mature AMs we analyzed AM development at E17.5 but observed no differences in frequency of immature (F4/80^int^ and CD11b^int^) AMs in WT and Siah1a/2-deleted cells (Figure 2A). CD11b and F4/80 expression levels on immature AMs were comparable in *cSiah1a/2*^*f/f*^*::Lyz2*^*Cre*^ and WT cells, but CD11c expression was significantly decreased in *cSiah1a/2*^*f/f*^*::Lyz2*^*Cre*^ immature AMs relative to WT cells, suggesting a differentiation delay (Figure 2B). At birth (day 0) the frequency of CD11b low and F480^+^ cells was comparable in WT and *cSiah1a/2*^*f/f*^*::CD11c*^*Cre*^ lungs (Figure 2C), but the frequency of CD11c^+^ SiglecF^+^ was markedly decreased in Siah1a/Siah2-deleted CD11b low F480^+^ cells relative to WT (Figure 2C, F). Within the population of Siah1a/2-deleted AMs (CD11b low F480^+^ SiglecF^+^ CD11c^+^ cells) we observed a significant decrease in CD11c and SiglecF expression relative to WT cells but observed no difference in F4/80 expression between genotypes (Figure 2D, F). We also observed an increased frequency of CD11c^-^ IMs and a decreased frequency of CD11c^+^ IMs (Figure 2E). These data suggest that Siah1a/Siah2 deletion does not change the frequency of fetal lung monocyte-derived pre-AMs but rather blocks their full maturation, based on decreased CD11c and SiglecF expression.

**Figure 2.**
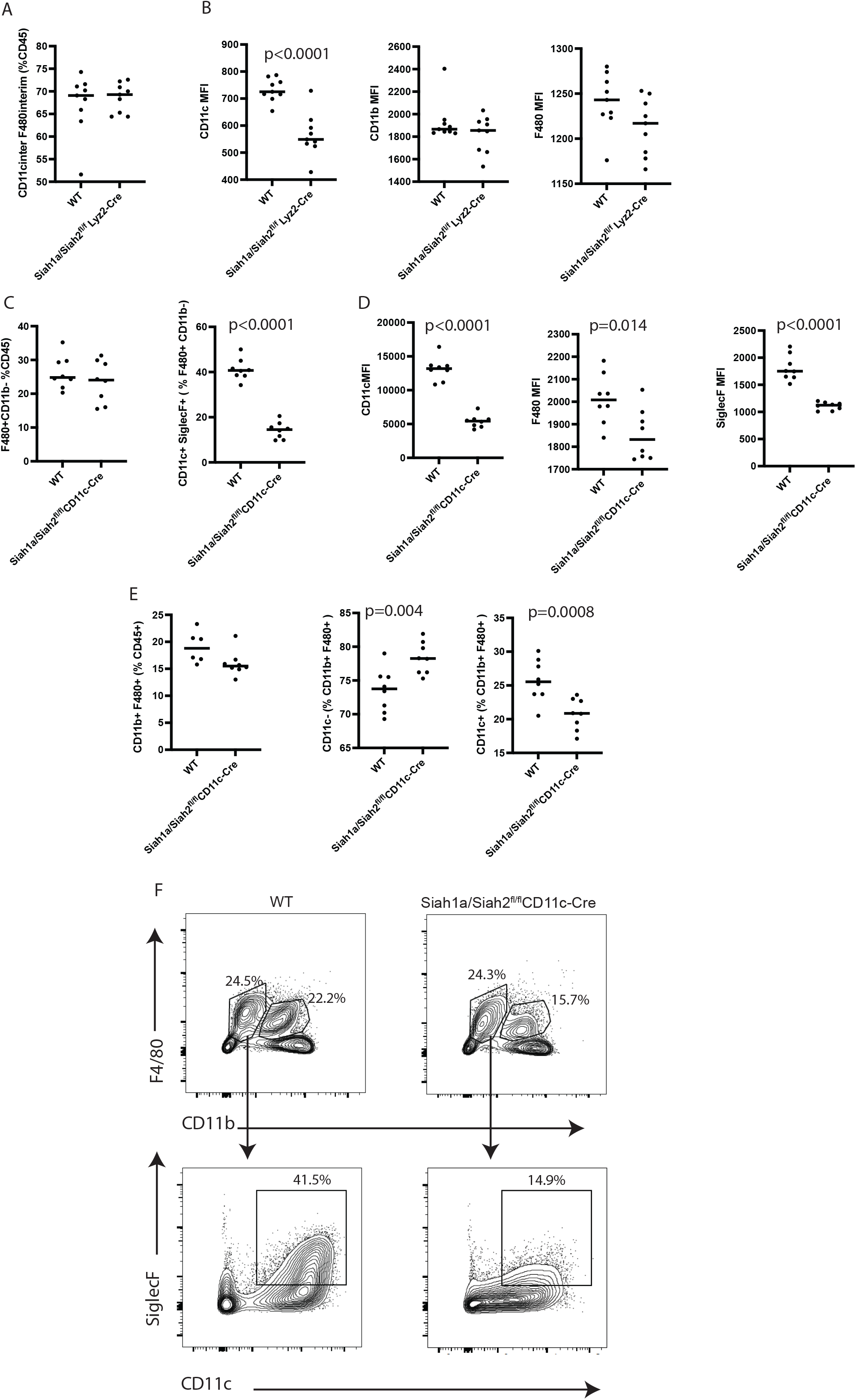
Siah1a and Siah2 regulate maturation of fetal monocytes to AMs. **A, B** Quantification of pre-alveolar macrophages showing intermediate expression of CD11c and F4/80 (A), and expression of CD11c, CD11b and F480 on Pre-AMs (B) from E17.5 lungs of *WT* and *cSiah1a/2*^*f/f*^*::Lyz2*^*Cre*^ embryos. Quantification is shown as the frequency of immune cells among CD45.2+ cells. n=9 for WT; n=9 for *cSiah1a/2*^*f/f*^*::Lyz2*^*Cre*^. **C**, Frequency of immune cells (F480^+^ CD11b low) in CD45^+^ cells, and of CD11c^+^ SiglecF^+^ immune cells in F480^+^ CD11b low immune cells from neonatal lungs (Day 0) of *WT* (n=8) and *cSiah1a/2*^*f/f*^*::CD11c*^*Cre*^ pups (n=8). **D**. CD11c, F480 and SiglecF expression on AMs (CD11b low F4/80+ SiglecF^+^ CD11c^+^) from neonatal lungs (Day 0) of *WT* (n=8) and *cSiah1a/2*^*f/f*^*::CD11c*^*Cre*^ pups (n=8). **E** Representative flow cytometry plots of the frequency of CD11c^+^ SiglecF^+^ cells by a gated subpopulation (CD11b low F480^+^ cells) of CD45.2^+^ cells from *WT* and *cSiah1a/2*^*f/f*^*::CD11c*^*Cre*^ lungs. n=8 for WT; n=8 for *cSiah1a/2*^*f/f*^*::Lyz2*^*Cre*^ Data were analyzed by unpaired t-test.

### AM regeneration from bone marrow requires Siah1a and Siah2

At steady state AMs self-renew while in inflammatory conditions or following bone marrow transplant, bone marrow-derived monocytes can replenish the AM population in the lung ^23,24^. To assess Siah1a/2 activity in bone marrow-derived AMs, we established a BM chimeric mouse model in which BM cells from *Lyz2*^*Cre*^ or *cSiah1a/2*^*f/f*^*::Lyz2*^*Cre*^ CD45.2^+^ mice were co-transferred with CD45.1^+^ cells into irradiated recipient mice. Then, 8 weeks later, we collected lungs from recipients and subjected cells to FACS analysis. Gating of mature AMs (CD11c^+^ SiglecF^+^ CD11b low and F480^+^) for CD45.1- and CD45.2-positive cells revealed that 70% of AMs were reconstituted with CD45.2^+^ WT cells, while only 50% of AMs were reconstituted with CD45.2^+^ and Siah1a/2-depleted cells (Figures 3A and 3B). Notably, CD11c and SiglecF expression significantly decreased within CD45.2^+^ Siah1a/2-depleted AMs, while of F4/80 expression was unchanged, suggesting inhibition of AM maturation (Figure 3C). Moreover, while 37% of CD11c^+^ IMs were reconstituted with CD45.2^+^ WT cells, ∼30% of CD11c^+^ IMs were reconstituted with CD45.2^+^ Siah1a/2-depleted cells (Figure 3D). Given that CD11c^-^ IMs and other immune cell components (monocytes, neutrophils, CD4 or CD8 cells) were equally represented in CD45.2^+^ WT and Siah1a/2-depleted cells (Figure 3E), we conclude that Siah1a/2 activity primarily impacts BM-derived AMs and CD11c^+^ IMs. As confirmation, we repeated the mixed bone marrow experiment using the *CD11c Cre* line. As in the *Lyz2 Cre* line, ∼70% of AMs were reconstituted with CD45.2^+^ WT cells, while only 30% of AMs were reconstituted with CD45.2^+^ Siah1a/2 depleted cells (Supp. Figure 2A). Within the CD45.2^+^ AM population CD11c expression significantly decreased in Siah1a/2-depleted relative to WT cells (Supp. Figure 2B), while we observed no differences in CD11c^+^ and CD11c^-^ IMs (Sup. Figure 2C). These data strongly suggest that Siah1a and Siah2 function in maturation of AMs from bone marrow-derived monocytes.

**Figure 3.**
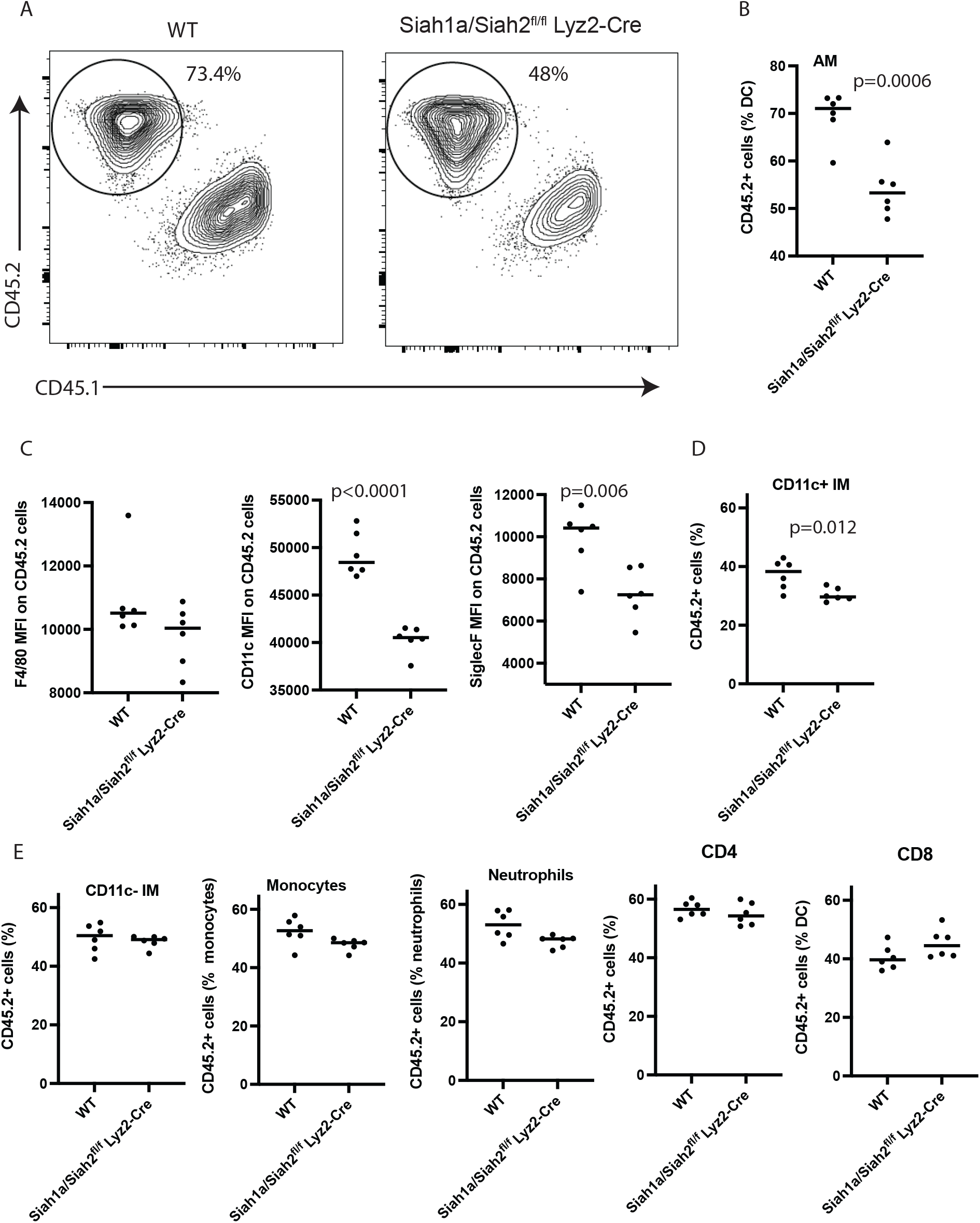
AM regeneration from in bone marrow requires Siah1a and Siah2. Bone marrow cells from *cSiah1a/2*^*f/f*^*::Lyz2*^*Cre*^ (CD45.2^+^) or CD45.2 WT mice were co-transferred with competitor CD45.1 bone marrow cells into lethally-irradiated C57Bl6 mice and lungs were harvested 10 weeks later. **A, B** Flow cytometry representation (A) and quantitation (B) of CD45.2-positive cells on lung AMs (CD11b low F4/80^+^ CD11c^+^ SiglecF^+^) from irradiated mice co-transfected with either a mix (1:1) of *cSiah1a/2*^*f/f*^*::Lyz2*^*Cre*^ CD45.2+ : CD45.1^+^ cells (n=6) or CD45.2 WT: CD45.1^+^ cells (n=6). **C** F4/80, CD11c, and SiglecF expression on CD45.2^+^ AMs of *cSiah1a/2*^*f/f*^*::Lyz2*^*Cre*^ CD45.2^+^ : CD45.1^+^ or CD45.2 WT: CD45.1^+^ co-transfected mice. n=6 for both genotypes. **D** Flow cytometry quantitation of CD45.2-positive cells on lung CD11c^+^ IMs (CD11b^+^ F4/80^+^ CD11c^+^) from irradiated mice co-transfected with either a mix (1:1) of *cSiah1a/2*^*f/f*^*::Lyz2*^*Cre*^ CD45.2^+^ : CD45.1^+^ cells (n=6) or CD45.2 WT: CD45.1^+^ cells (n=6). **E** Flow cytometry quantitation of CD45.2-positive cells on lung CD11c^-^ IMs (CD11b^+^ F4/80^+^ CD11c^-^), monocytes (CD11b^+^ Ly6c), neutrophils (CD11b^+^ Ly6G^+^), and CD4^+^ or CD8^+^ cells from irradiated mice co-transfected with either a mix (1:1) of *cSiah1a/2*^*f/f*^*::Lyz2*^*Cre*^ CD45.2+ : CD45.1+ cells (n=6) or CD45.2 WT: CD45.1+ cells (n=6). Data were analyzed by unpaired t-test.

### Siah1a/2 controls immunoregulatory function of AMs under homeostatic conditions

To assess changes in gene expression and signaling pathways in AMs following Siah1a/2 depletion we performed RNA-seq analysis of lung cells (n=3 for WT and KO, n=1 for Cd11c three lungs per sample) sorted by FACS based on CD11c^+^ F480^+^ SiglecF^+^ expression from WT, *cSiah1a/2*^*f/f*^*::Lyz2*^*Cre*^ and *cSiah1a/2*^*f/f*^*::CD11c*^*Cre*^ mice. Principal components analysis showed a similar gene expression patterns of samples from *cSiah1a/2*^*f/f*^*::Lyz2*^*Cre*^ and *cSiah1a/2*^*f/f*^*::CD11c*^*Cre*^ mice, as well as a distinct segregation between WT and *Siah1a/2*-depleted macrophages along main component PC1 (Figure 4A). Differential gene expression analysis confirmed similar gene expression signatures in AMs from *cSiah1a/2*^*f/f*^*::Lyz2*^*Cre*^ and *cSiah1a/2*^*f/f*^*::CD11c*^*Cre*^ mice and a reduced expression of Siah1a and Siah2 in Siah1a/2-deleted AMs (Figure 4B). Relative to WT AMs, we identified 565 upregulated and 363 downregulated genes distinctly expressed in AM samples from *cSiah1a/2*^*f/f*^*::Lyz2*^*Cre*^ mice (Figure 4B, Sup Figure 3A). Given both the immature-like state of AMs from Siah1a/2-depleted MΘ and the importance of GMCSF/PPARγ signaling in AM differentiation and maturation, we asked whether components of these pathways (namely, GMCSF (Csf2), Tgfβ1, Bach2, Tgfbr1, Tgfbr2, Csf2ra, and Csf2rb) were differentially expressed. Except for a small but significant increase in Csf2rb and Tgfbr2 expression seen in Siah1a/2-depleted macrophages, expression of all other GMCSF components or their downstream effectors (PPARγ) was comparable among genotypes (Sup. Figure 3B).

**Figure 4.**
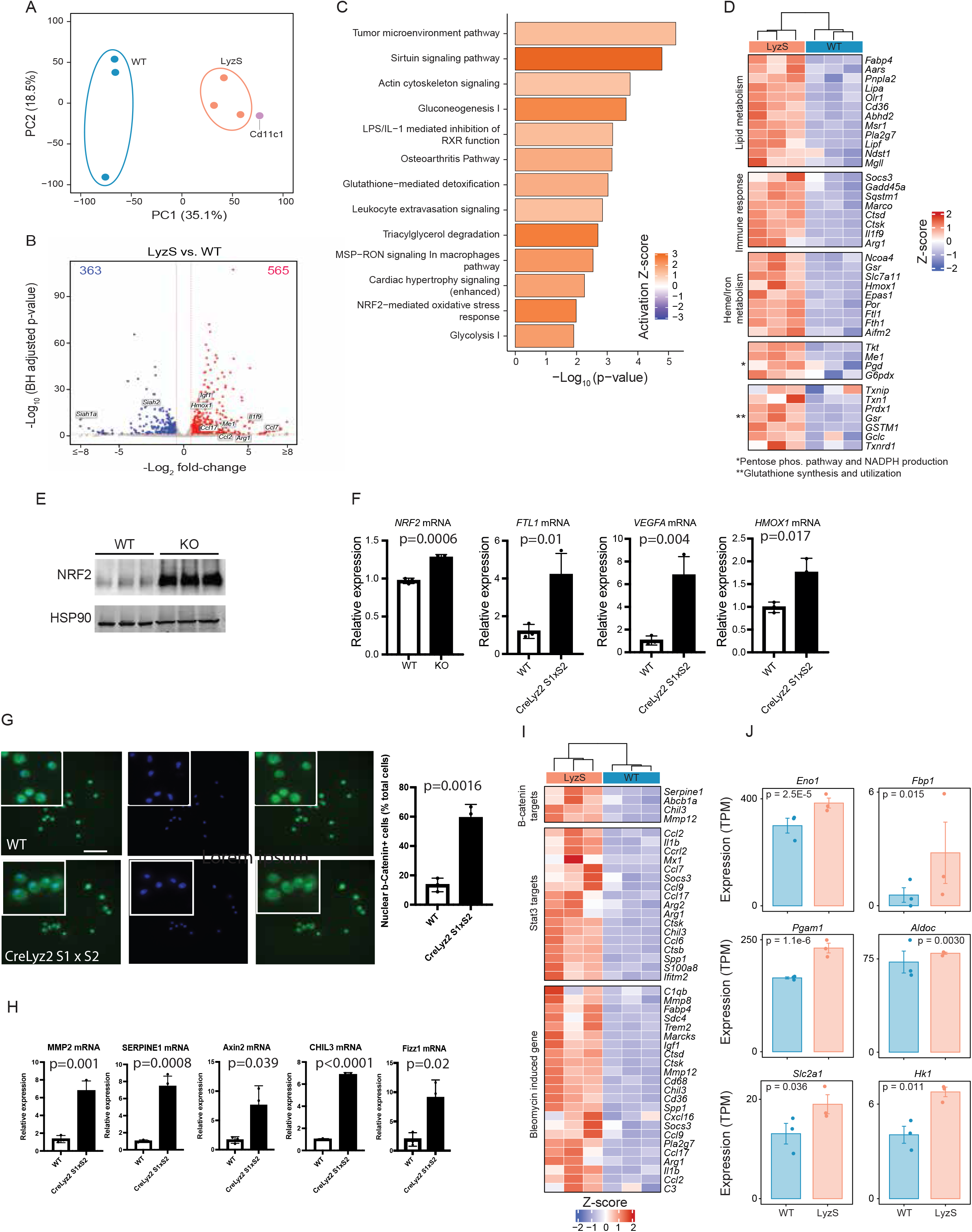
Siah1a and Siah2 control AM immunoregulatory function in homeostatic conditions. RNA-seq analysis of CD11c+ F480+ SiglecF+ cells sorted from lungs of WT (n=3), *cSiah1a/2*^*f/f*^*::Lyz2*^*Cre*^ (n=3), or *cSiah1a/2*^*f/f*^*::CD11c*^*Cre*^ (n=1) mice. Each sample was a pool of 3 lungs. **A** Principal component analysis (PCA) plot based on gene expression. WT and KO samples segregate along the main principal component (PC1). **B** Volcano plot of genes differentially expressed in KO versus WT lung, based on RNA-seq. Selected up- and down-regulated genes are shown. N= 3 for both genotypes. **C** Bar plots depicting the activation z-scores and significance p-values of the most upregulated pathways in *cSiah1a/2*^*f/f*^*::Lyz2*^*Cre*^ *compared to WT* AMs. **D** Heatmap showing row-wise z-scores of expression of selected genes associated with particular functions induced by NRF2 from WT and *cSiah1a/2*^*f/f*^*::Lyz2*^*Cre*^ AMs based on RNA-seq analysis. n = 3 for both genotypes. Samples (columns) were hierarchically clustered. **E** Primary AMs were collected after 6 days of culture for western blot analysis. WT and *cSiah1a/2*^*f/f*^*::Lyz2*^*Cre*^ AM lysates were immunoblotted for NRF2. HSP90 was used as a loading control. n = 3 for both genotypes. **F** Primary AMs were cultured 6 days for QPCR analysis of indicated transcripts. n = 3 for both genotypes. **G** (left) β-catenin staining of primary AMs cultured overnight in media containing GM-CSF and (right) quantification. n =3 for each genotype. β-catenin = green, Dapi = blue. Scale bar, 600 µM. **H** Primary AMs were cultured 6 days for QPCR analysis of indicated transcripts. n = 3 for both genotypes. **I, J** Expression of transcripts induced by β-catenin/STAT3 or following treatment with bleomycin (I) implicated in glycolysis (J) from WT and *cSiah1a/2*^*f/f*^*::Lyz2*^*Cre*^ AMs based on RNA-seq analysis. n = 3 for both genotypes. Data in panel F, G and H were analyzed by unpaired t-test, data in panel J were analyzed by Wade test.

To identify pathways deregulated by Siah1a/2 ablation, we performed pathway analysis using IPA of the above differentially expressed gene. Among the most activated signaling pathways were tumor microenvironment, actin cytoskeleton and NRF2 (Figure 4C). NRF2 is tightly regulated at transcriptional and post-translational levels, with its key guardian, KEAP1, controlling its stability ^25, 26, 27^. Notably, under hypoxic or low glucose conditions Siah2 has been implicated in control of NRF2 stability ^28, 29^. Accordingly, we observed induction of NRF2-target genes, including those involved in diverse metabolic activities, NADPH production, and the immune response, in samples from Siah1a/2-deficient relative to WT AMs (Figure 4D). Moreover, Western blot analysis of primary AM cultures from lungs of these mice, confirmed a marked increase in levels of NRF2 protein (Fig 4E) (but not mRNA (Fig 4F) and a corresponding increase in mRNAs encoding NRF2 targets, including Ftl1, Hmox1, and Vegfa in *cSiah1a/2*^*f/f*^*::Lyz2*^*Cre*^ AMs when compared to WT (Figure 4F). At the same time, immunofluorescence revealed comparable nuclear localization of PPARγ in AMs from both genotypes, excluding relevance of the PPARγ pathway to Siah1a/2 activity in this context (Sup. Figure 3C). Of note, differences in NRF2 levels and those of its target genes were restricted to AMs and not seen in BM-derived macrophages (Sup figure 3D).

We next addressed mechanisms underlying upregulation of cytoskeletal signaling seen in Siah1a/2-deficient AMs (Figure 4C). Among Siah2 substrates that could be implicated in its control of cytoskeletal organization and cell migration is β-catenin ^30, 31, 32^. β-catenin is also implicated in AM differentiation ^33^, as it drives immature-like phenotypes similar to the one observed in Siah1a/2-ablated AMs. Indeed, isolated AMs lacking Siah1a/2 showed increased nuclear localization of β-catenin, indicative of activation (Figure 4G). Moreover, immunohistochemistry with an antibody specific for the active form of β -catenin showed more nuclear and cytoplasmic intense staining in primary cultures of Siah1a/2-deficient compared to WT AMs (Supp Figure 3E). Activation of beta catenin in macrophages was found to potentiate STAT3 activity ^34, 35, 36^. Accordingly, qPCR analysis of primary AM cultures confirmed a marked increase in expression of β-catenin targets and STAT3 targets, including Axin2, Serpine1, Mmp2, Fizz1 and Chil3 (Figure 4H). Likewise, IPA analysis of RNA-seq data revealed increased transcription of β-catenin- and of STAT3-activated genes, including metabolic signaling factors (glycolysis) linked with β -catenin activation, in Siah1a/2-ablated AMs that are maintained under a homeostatic state when compared to WT (Figure 4I, J). Independent support for a link between Siah1a/2 and a unique inflammatory signature observed in AMs comes from the identification of a gene signature reported in lungs of mice subjected to bleomycin treatment, a chemical widely used to induce lung fibrosis (Figure 4I). Of note, during bleomycin induced fibrosis β-catenin activity is increased in alveolar macrophages ^37^ and mice with conditional β-catenin knockdown driven by the CD11c promoter show improvement in fibrosis after bleomycin treatment ^33^. Overall, these data reveal that Siah1a/2 deletion in AMs increases NRF2 and beta catenin signaling, and that Siah1a/2-deleted AMs display a profibrotic immune signature associated with tumor progression (Figure 4 D and I).

### Siah1a and Siah2 deletion in macrophages promotes urethane-induced lung cancer

Given AM abundance in normal lung, we asked whether Siah1a/2 expression in AMs alters the course of lung cancer by subjecting mice harboring either WT or Siah1a/2-deficient AMs to treatment with urethane, a carcinogen that induces KRAS mutations and promotes lung tumor development ^38^. Lungs were collected from mice 24 weeks after the first of six weekly urethane injections. Notably, mice lacking Siah1a/2 in AMs showed increased numbers of tumor nodules (Figure 5A) and increased tumor size (Figure 5B), based on comprehensive assessment of serial lung sections (Figure 5C). While the frequency of CD45^+^ cells was comparable between genotypes (Supp. Figure 4A), AM frequency was lower in tumor-bearing mice (12% of CD45^+^ cells, Figure 5D) compared to tumor free mice (22% of CD45^+^ cells Figure 1C), with the most notable decrease observed in tumor bearing Siah1/2 deleted AMs (4% of CD45^+^ cells, Figure 5D). Of note, immature-like AMs, not observed in WT tumor free mice (Figure 1E, F), were observed in WT after urethane administration, with the greatest increase seen among Siah1a/2-ablated AMs (Figure 5D, 5E, 5F). Among mature (SiglecF^+^ CD11c^+^ F480^+^ CD11b^-^) AMs population, Siah1a/2-deleted AMs exhibited a significant decrease in expression of CD11c (67%), SiglecF (36%) and the co-stimulatory protein CD80 (80%) relative to WT after urethane administration, while the expression of CD206 and MHCI was comparable (Figure 5G, Sup. Figure 4B). We also observed a small but significant increase in the number of CD11c^-^ IMs and a decrease in CD11C^+^ cells (Figure 5H and I), as seen in tumor-free mice. Siah1a/2-deficient IMs (Lyz^Cre^) exhibited comparable levels of MHCI, CD206, CD11c and CD80 within Cd11c^+^ IMs to that seen in WT cells (Sup Figure 4C), suggesting that both genotypes exhibit similar degree of activation despite the lower frequency of CD11c^+^ IMs seen after Siah1a/2 depletion. While there were no differences in the number of B cells or CD4 and CD8 cells, we observed a marked decrease in CD44^+^ cells within the CD8 population, along with a lower frequency of NK1-positive and active (B220^+^ among NK1.1^+^) NK1^+^ cells after Siah1a/2 ablation (*Lyz2*^*Cre*^) relative to WT cells (Sup. Figure 4D).

**Figure 5.**
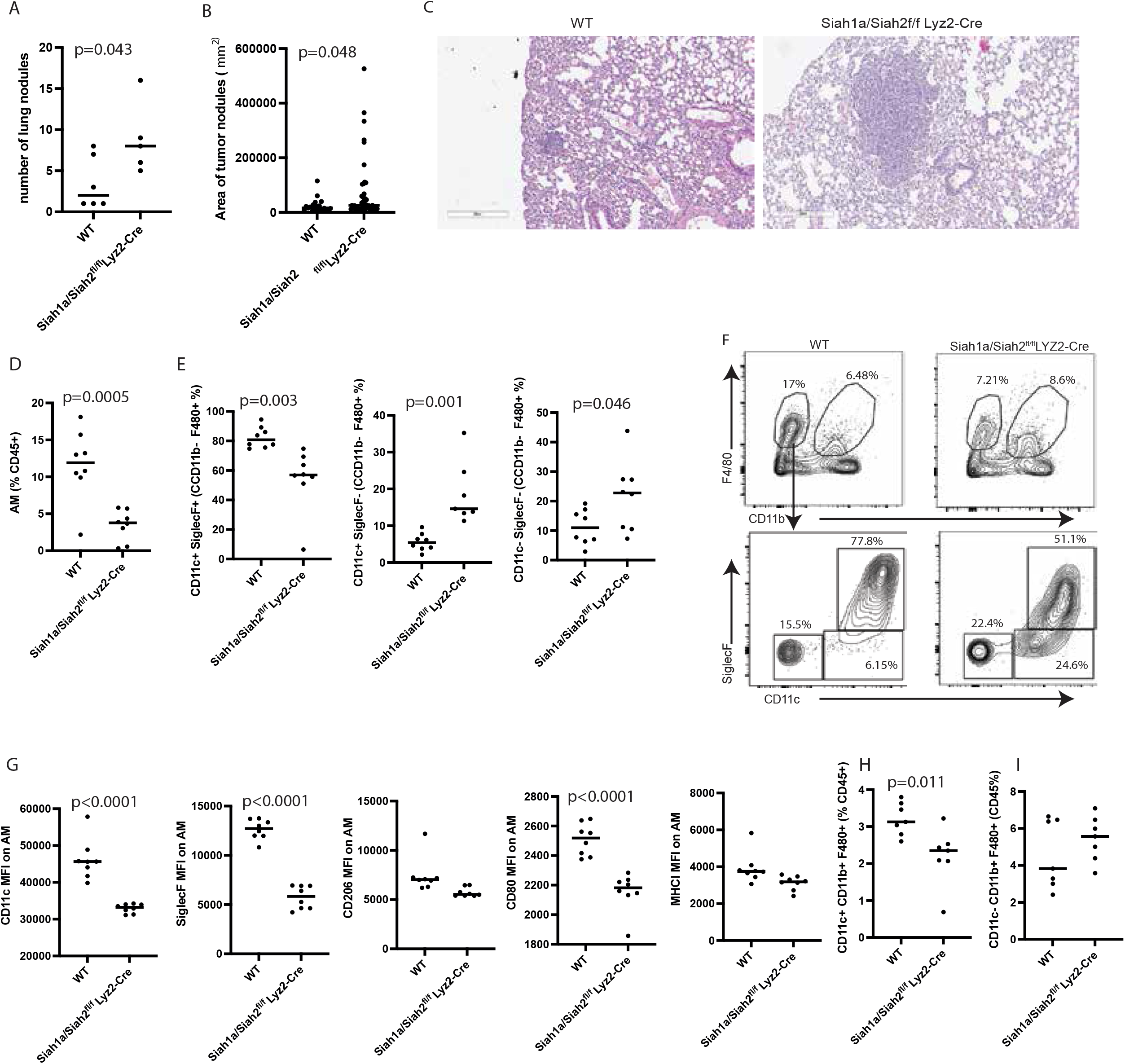
Siah1a and Siah2 deletion in macrophages promotes lung cancer following urethane treatment. Urethane (1mg/g mouse weight) was injected intraperitoneally once a week for 6 weeks and lungs were collected 22 weeks after the first injection. **A, B** Lungs were processed, and lung nodule number (A) and size (B) were quantified by H&E staining. n=6 for WT, n=5 for *cSiah1a/2*^*f/f*^*::Lyz2*^*Cre*^ in panel A; n=21 for WT, n=44 for *cSiah1a/2*^*f/f*^*::Lyz2*^*Cre*^ in panel B. **C** Representative image of H&E staining of lungs for each genotype. Scale bar, 300 µM. **D** AM frequencies among CD45+ lung cells after urethane injection. n = 8 for each genotype. **E** Frequencies of CD11c^+^ SiglecF^+^ (mature macrophages) and CD11c^+^ SiglecF^-^, CD11c^-^ SiglecF^-^ (immature AMs) among CD11b low F4/80^+^ cells. n = 8 for each genotype. **F** Representative flow cytometry plots showing CD11c^+^ SiglecF^+^ expression by a gated subpopulation (CD11b low F480^+^ cells) of CD45.2^+^ cells from *WT* and *cSiah1a/2*^*f/f*^*::Lyz2*^*Cre*^ lungs. **G** Expression of CD11c, SiglecF, CD206, CD80 and MHCI on an AM (CD11b low F4/80^+^ CD11c^+^ SiglecF^+^) population. n = 8 for each genotype. **H, I** Frequencies of CD11c^+^ (H) and CD11c-(I) IMs among CD45+ cells. n = 8 for each genotype. Data were analyzed by unpaired t-test.

### Siah1a/2 deletion in macrophages promotes a pro-fibrotic phenotype

To identify signaling pathways in immune cells altered by Siah1a/2 deletion in lung macrophages, we performed single cell RNAseq (scRNA-seq) of CD45^+^ cells enriched from lungs collected from WT (n=1, pool of 3 lungs) and tumor-bearing mice (n=1, pool of 3 lungs) at the end of 6 weeks of urethane treatment. Integrated analysis of WT (n=7884 cells) and KO (n=6035 cells) scRNA-seq datasets identified 17 immune cell clusters with distinct gene expression patterns (Figure 6A and 6B). Of these 17 clusters, clusters 1 and 13 were positive for Ncr1 and Klrb1a, markers of NK cells; cluster 12 was positive for the neutrophil marker Csf3r; cluster 15 expressed B cell markers Cd19 and Pax5; and clusters 0, 4, 6, 7, 8 and 10 expressed the T cell marker Cd3e. The myeloid population expressing Csf1r was represented by clusters 2, 3, 5 and 11. We identified plasmacytoid dendritic cells (DCs) in cluster 9 (Cd209a, Itgax, H2-aa), while DCs were also identified in cluster 16 (H2-aa, Flt3 and Ly75), and basophils (Fcer1a and Cd200r3) in cluster 14 (Figure 6B).

**Figure 6.**
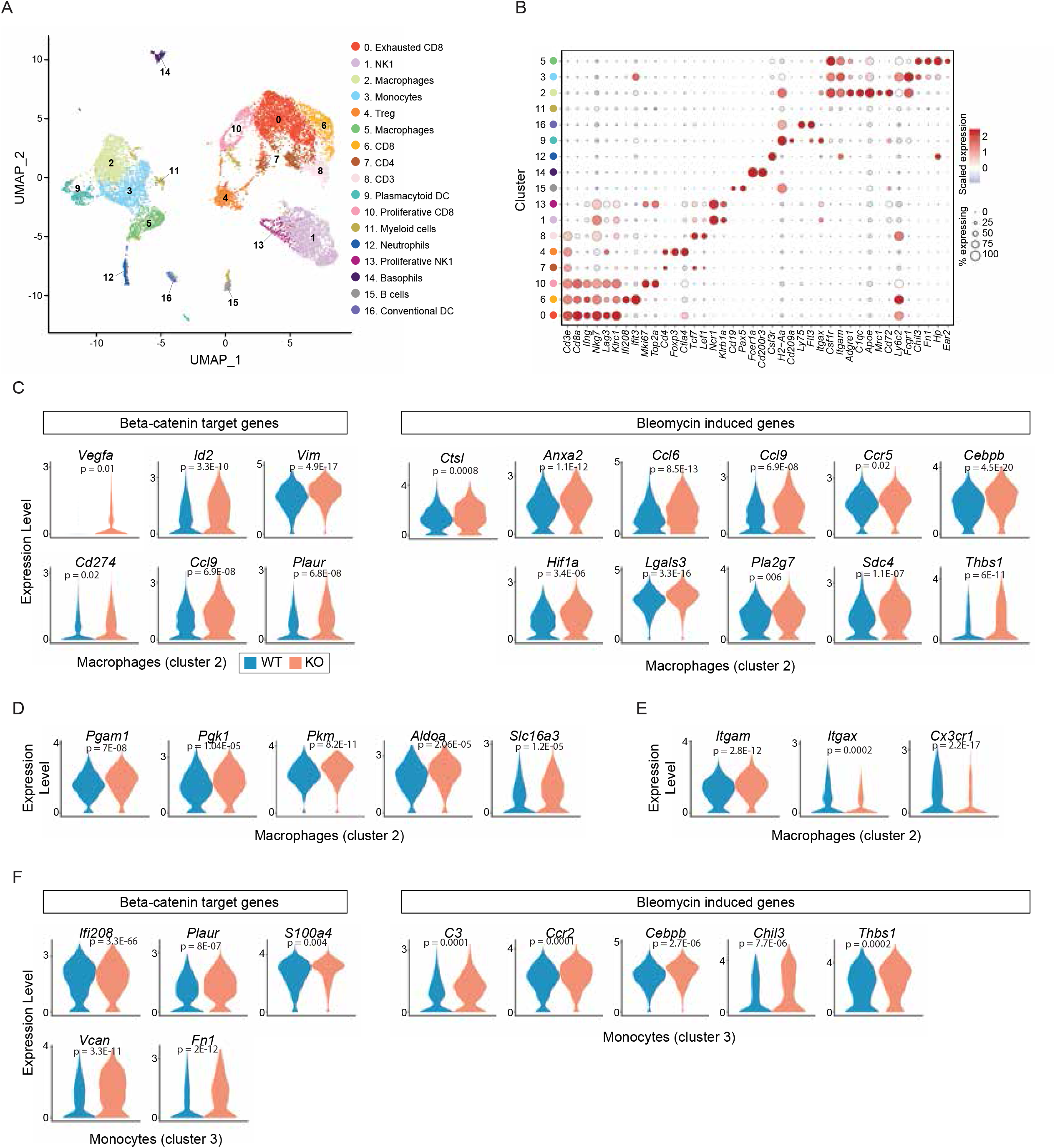
Siah1a and Siah2 deletion in macrophages promotes an immature state and induces an immunoregulatory/pro-fibrotic phenotype in specific macrophage clusters. CD45^+^ cells were sorted by flow cytometry from WT and *cSiah1a/2*^*f/f*^*::Lyz2*^*Cre*^ lungs collected 7 weeks after the first urethane injection, and single cell RNA-seq was performed. **A** UMAP plot of CD45^+^ cells from WT (n=7884 cells) and *cSiah1a/2*^*f/f*^*::Lyz2*^*Cre*^ (n=6035 cells) lungs showing 17 distinct clusters. Immune cell types identified using expression of specific cell markers are labeled. **B** Scaled expression levels of selected markers used to identify clusters in the UMAP of CD45+ cells. **C-E** Violin plots comparing expression of genes in WT and *cSiah1a/2*^*f/f*^*::Lyz2*^*Cre*^ cells in macrophages (cluster 2) regulated by β-catenin or modulated upon treatment with bleomycin (C) or functioning in glycolysis (D) or macrophage differentiation (E). **F** Violin plots comparing expression of genes in monocytes (cluster 3) regulated by β-catenin or modulated by bleomycin treatment. Data were analyzed using *FindMarkers()* and *MAST* test in Seurat.

Within T cell populations, CD8^+^ cells were distributed in 3 clusters: cluster 0, which included terminally effector exhausted CD8^+^ cells expressing genes involved in anti-tumor immunity like Ifnγ and Nkg7, and genes with immunosuppression function like Lag3, Klrc1, and Ctla4; cluster 6, consisting of CD8^+^ effector T cells expressing Ifnγ and the interferon response genes Ifi208 and Ifit3; and cluster 10, representing proliferating effector CD8^+^ cells expressing Mki67 and Top2a. Among CD4^+^ populations, cluster 7 consisted of CD4^+^ effector cells producing Ifnγ? while cluster 4 represented Foxp3-expressing CD4^+^ T regulatory cells. Cluster 8 included T cells negative for CD4 and CD8 and expressing Tcf7 and Lef1 (Figure 6B).

Changes in immune cell frequencies in Siah1a/2-ablated AMs included a notable increase in neutrophils represented in cluster 12 (2.45% in *cSiah1a/2*^*f/f*^*::Lyz2*^*Cre*^ cells vs 1.15% in WT cells) concomitant with a decrease in CD8 effector/proliferative cells represented in cluster 10 (2.93% in *cSiah1a/2*^*f/f*^*::Lyz2*^*Cre*^ cells *vs*. 4.46% of WT cells, Sup. Figure 5A). Within the myeloid population of Csf1r^+^ cells (clusters 2, 3, and 5), cluster 2 exhibited the highest expression of Adgre1 (F4/80) and was enriched of cells positive for AM markers including Apoe, Mrc1 (CD206), Cd72 and Itgax (Figure 6B). This cluster was also positive for genes indicative of monocytic origin (Ly6c2, Fcgr1 and H2-a) and may represent monocytes differentiating into AMs (Figure 6B). Cells in cluster 3 expressed highest levels of Ly6c, indicative of monocytes, while cluster 5 was represented by macrophages expressing Hp, Ear2, Ace, Chil3 and Fn1 (Figure 6B).

Among upstream regulators, IPA analysis identified significant upregulation of β-catenin signature genes and bleomycin-induced genes (profibrotic genes) (Figure 6C) in cluster 2, which contained residential and monocyte-derived AM populations. Among these, were genes known to be involved in tumor progression, (Ccl6, Ccl9, Vegfa, Anxa2 and Lgals3) (Figure 6C). Siah1a/2-deleted cells of cluster 2 also showed increased expression of glycolysis-associated genes relative to WT, suggestive of increased of macrophage activity and associated to a pro-fibrotic phenotype in AMs^39^ (Figure 6D). Differential gene expression of cluster 5 exhibited increased mitochondrial respiration in Siah1a/2 deleted cells and an increased expression of Fn1 (Sup. Figure 5B). Macrophages possessing an immature-like phenotype were enriched upon Siah1a/2 ablation in cluster 2, as reflected by increased Itgam (CD11b) expression concomitant with decreased Itgax (CD11c) and Cx3cr1 expression (Figure 6E). Notably, Siah1/2-deleted monocytes (cluster 3) exhibited increased expression of β-catenin-activated genes (i.e., Plaur, Fn1 and S100a4) and profibrotic genes (bleomycin induced genes), which are implicated in lung fibrosis (Figure 6F).

Although we observed increased NRF2 activity in tumor free lungs, as evidenced by Hmox1, Ftl1 or Fth1 upregulation in Siah1a/2-ablated WT AMs, we did not observe these increases after urethane treatment, likely since urethane induces an oxidative /inflammatory phenotype that may mask effects of Siah1a/2-depletion (Sup Figure 5C). Consistently, U-map analysis revealed high expression of NRF2 in both WT and Siah1a/2 deleted myeloid population (Sup. Figure 5C) indicative of urethane-induced oxidative stress ^40, 41^ which led to increased NRF2 expression and activity. Of note, the inflammatory signature seen in Siah1a/2-deficient AMs in tumor-free lungs was maintained in tumor-bearing lungs (Figure 6C). Accordingly, AMs presenting an immature-like state reportedly exhibit a unique inflammatory signature in pathologies similar to that seen in fibrotic macrophages ^23, 42, 43^.

Initial analyses of the 3 myeloid clusters (2, 3, 5) did not separate residential AMs from monocytes derived AMs. Re-clustering of those 3 groups (Figure 6A) gave rise to 9 myeloid subclusters (Figure 7A). All expressed genes associated with monocytic origin (Ccr2, Cx3cr1, Ly6c2, H2-aa and H2-ab1; Figure 7B). Among the 8 subclusters, 1, 2, and 3 exhibited highest Ly6c2 and Sell expression, a characteristic of monocyte populations. Subcluster 3 also exhibited highest expression of Hp and Fn1. Subclusters 5, 6, 7, and 8 showed lowest Ly6c2 and Ccr2 expression, indicating they contain more differentiated residential macrophages (Figure 7C). Subclusters 5 and 8 exhibited AM markers (Itgax, Mrc1, Apoe and Siglec1), reflective of a mature state. By contrast, subcluster 4 was CD3e-positive, indicating a unique group of T cells also positive for F4/80 and CD11c (Figure 7C). Lastly, subcluster 0 was positive for markers of monocytic origin (Ly6c2, Ccr2 and H2-aa) and for AM markers (Itgax, Mrc1, Apoe, Cd74, Cxcl16 and Cd72, which were also found in subcluster 5), suggesting that cluster 0 contains immature AM of monocytic origin. Of note, while subclusters 5 and 8 showed highest expression of immunosuppressors (C1qa, C1qb and C1qc), subcluster 0 showed intermediate levels of those genes, confirming the transition state of cluster 0 cells (Figure 7C).

**Figure 7.**
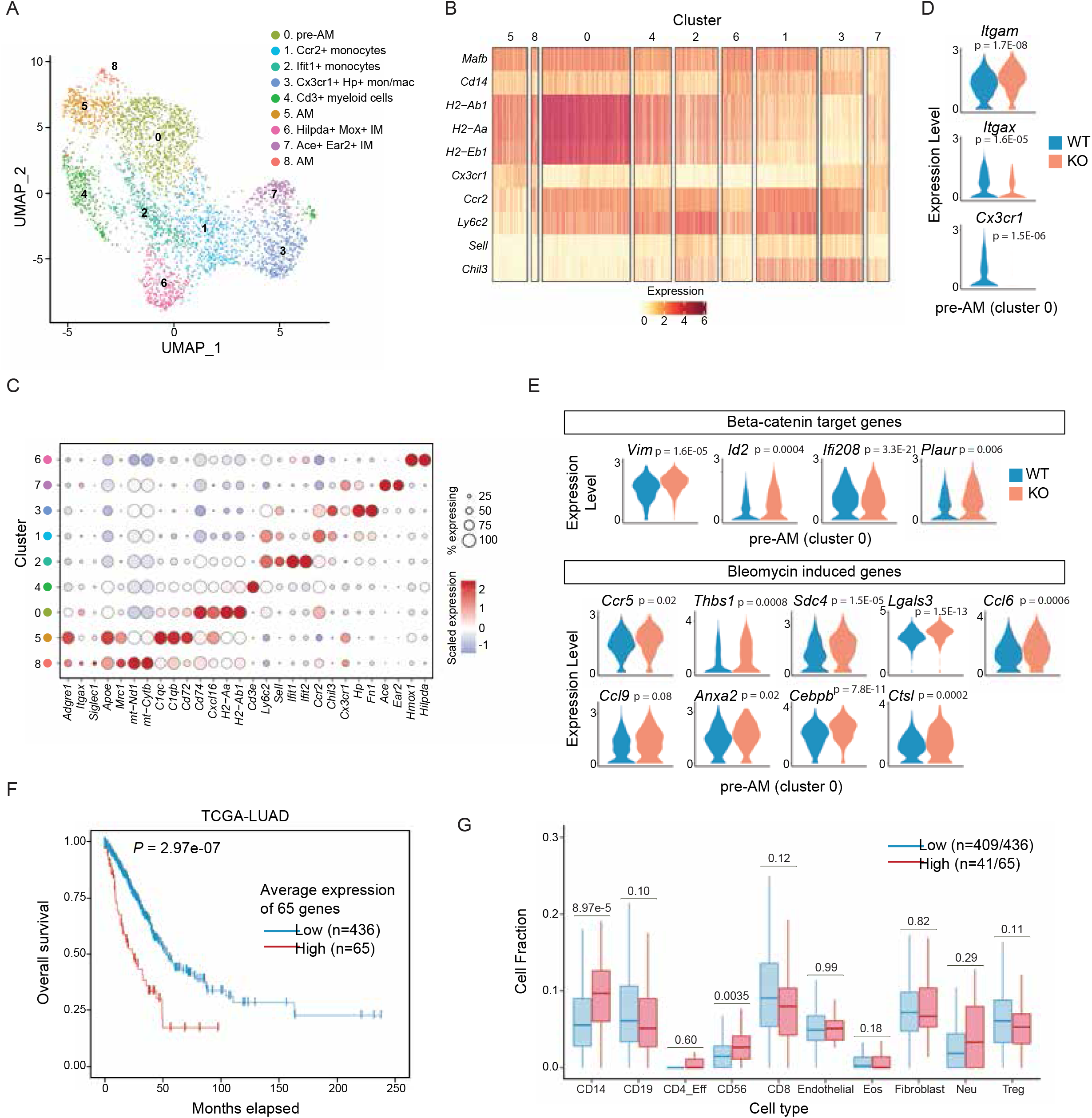
Siah1a and Siah2 deletion in macrophages promotes an immunoregulatory/pro-fibrotic phenotype in monocyte-derived AMs. Myeloid cell clusters (2, 3, and 5 from global UMAP in Figure 6A) from single cell RNA-seq analysis of CD45^+^ cells were re-clustered to obtain higher resolution clustering of myeloid cells. **A**. UMAP plot of myeloid cells from WT (n=2245 cells) and *cSiah1a/2*^*f/f*^*::Lyz2*^*Cre*^ (n=1417 cells) lungs showing 9 distinct clusters. **B**. Heat map of expression levels of markers of monocytic origin in each myeloid cell cluster. **C**. Dot plots showing scaled expression of marker genes used to identify specific sub-clusters of myeloid cells. **D, E**. Violin plots comparing expression of genes from sub-cluster 0 functioning in AM differentiation (D) or regulated by β-catenin or following bleomycin treatment (E) in each genotype. **F** Average expression of 65 out of 72 upregulated genes in sub-cluster 0 (KO vs. WT) found in TCGA-LUAD RNA-seq dataset was used to stratify patients as high (n=65) and low (n=436) expressors of this 65-gene signature. Patients with high expression of the gene signature show worse overall survival (p-value=2.97e-07, log rank test) compared to low expressors. **G** Cell type compositions of TCGA-LUAD patient tumors were assessed using cell-type and gene-expression deconvolution dataset from CODEFACS. The same patient stratifications as Figure 7F were used. Shown are comparisons of 10 cell type composition fractions of high and low expressors of the inflammatory gene signature in sub-cluster 0. P-values for composition difference in each type cell type are shown (Wilcoxon rank sum test). Data in panels D and E were analyzed using *FindMarkers()* and *MAST* test in Seurat.

Decreased proportions of AMs in subclusters 5, among CD45^+^ cells from lungs of Siah1a/2-depleted macrophages (Supp Figure 6A) compared to WT, confirmed our earlier FACS analyses. Likewise, the immature-like state of AM in cluster 0 was confirmed by gene expression analysis showing increased Itgam and decreased Itgax and Cxc3r1 expression in Sia1a/2-depleted macrophages (Figure 7D). Lastly, increased expression of genes implicated in glycolysis (Supp. Figure 6B), a metabolic reprogramming found in pro-fibrotic AMs^39^, along with increased β-catenin signaling and bleomycin induced genes was confirmed in Siah1a/2-depleted macrophages (Figure 7E). These data indicate that Siah1a/2 loss in AM promotes pro-tumorigenic and pro-fibrotic phenotypes. Further analysis excluded the possibility that the immature-like state of Siah1a/2-deleted AMs was due to dysregulated recruitment of bone marrow-derived monocytes. Specifically, LY6C protein expression in AMs, immature AMs and IMs (Sup. Figure 6C) was comparable in WT and Siah1a/2-mutant cells, although we noted differences between IM and AM populations: LY6C was highest in CD11b^-^ F480^+^ CD11c^-^ and SiglecF^-^ cells (immature AMs) and in CD11c^-^ IMs and lowest in mature AMs (10%). These data suggest that immature-like phenotypes seen in Siah1a/2-deleted AMs result from cell-autonomous effects rather than from impaired lung infiltration by monocytes.

To assess the possible clinical relevance of these findings, we determined the expression of genes that were significantly upregulated (65 genes) in cluster 0 (immature AMs) of Siah1a/2 deleted cells (compared to WT; Supplemental Table 1) in two independent data sets. First, the expression of these 65 genes in TCGA database of LUAD (lung adenocarcinoma) cohort identified patients with poor prognosis and shorter survival (Figure 7F). Second, using a precomputed estimates of cell abundance or cell-type-specific signature based on deconvolving cell-type-specific gene expression in each sample from bulk expression^44^ of LUAD patients, patients with higher average expression of the unique signature (65 genes) identified in Siah1a/2 ablated macrophages were also found to have increased myeloid cells (CD14^+^) infiltration (Figure 7G). The TCGA dataset of the LUAD was also used to identify which of the 65 genes are associated with worse survival. To this end, we calculated the hazard ratios for each of the 65 inflammatory genes that were significantly upregulated in cluster 0 (immature AMs of Siah1a/2 mutant versus WT cells) identifying genes involved in glycolysis (i.e., lactate production and export *LDHA, SLC16A3*), in generation of plasmin and degradation of extracellular matrix (*PLAUR, S100A10/ANXA2, CTSL, ADAM8*) (Sup. Figure 7D). Of note, these signaling pathways have been observed in pro-fibrotic macrophages and implicated in cancer progression. Overall, these data establish the relevance of our finding in the urethane-induced lung cancer mouse model to human tumors.

## Discussion

We report here that Siah1a and Siah2 loss in AMs promotes their immature-like state and emergence of a profibrotic gene signature. Moreover, we show that Siah1a and Siah2 activity likely enables alveolar macrophage differentiation and thus impacts lung inflammation. Selective ablation of Siah1a and Siah2 in macrophages was performed using two complementing genetic approaches: the Lyz2-Cre and the CD11c-Cre models. While each system shows some cell type selectivity, both are expressed in AMs, supporting our conclusion that inflammatory phenotypes seen after Siah1a/2 ablation by either system are primarily due to AMs activity. Using both approaches we define the role of Siah1a and Siah2 in the shift of AMs from mature to an immature and pro-inflammatory phenotype, which was most pronounced following lung carcinogen administration. Mechanistically, we show that two substrates destabilized by Siah1a/2, NRF2 and β-catenin, mediate pro-tumorigenic phenotypes, suggesting that Siah1a/2 ubiquitin ligases prevent inflammatory and profibrotic signaling by limiting NRF2 and β-catenin availability and activity. Notably, Siah1a/2 effects were mainly restricted to AMs, the most abundant population of lung macrophages. While AMs reportedly function in turnover of pulmonary superfactants and removal of dead cells from alveoli, their differentiation is thought to be primarily regulated by GM-CSF/PPARγ and TGFβ signaling ^16, 45^. Here we show that Siah1a/2 contribute to AM maturation independently of the GM-CSF/PPARγ pathways, revealing an unexpected signaling path underlying AM maturation.

NRF2 controls central cellular homeostatic pathways regulating reactive oxygen species, cellular redox, ferroptosis, apoptosis and key metabolic pathways ^45^. Notably, NRF2 undergoes extensive post-translational modifications that dictate its stability and activity, primarily by KEAP1. Regulation by Siah1a/2 represents a novel layer of NRF2 regulation likely in response to stress signaling, such as oxygen or glucose deprivation, activities often associated with Siah1a/2 function. Here we show that increased NRF2 protein level induces a set of target genes that are implicated in lung cancer progression, namely Arg1, VEGFA and HMOX1. The increased expression of these genes and others are conducive to cancer cells growth and proliferation.

β-catenin signaling reportedly modulates the immune response in macrophages, in part through STAT3-dependent IL4 signaling ^34, 35, 36^. In hepatocellular carcinoma, β-catenin activation in TAMs is linked with macrophage polarization to an M2 type ^36^. Likewise, increased β-catenin levels in AMs have been observed following treatment with the chemotherapy drug bleomycin ^37^, which promotes fibrosis. Notably, fibrosis is among the pathological condition enriched with immature-like alveolar macrophages ^23, 24, 42^.

Our findings in a lung carcinogen model confirm that Siah1a/2 controls AM differentiation and inflammatory signaling. Urethane induces pulmonary adenocarcinoma resembling human adenocarcinoma with KRAS mutations ^38^. We observed increased numbers of larger lung tumors in mice lacking Siah1a/2 in macrophages relative to controls. Siah1a/2 ablation increased the number of exhausted T cells and decreased the number of activated NK cells in lung tumors concomitant with increased expression of genes implicated in tumor proliferation and immunosuppression. Notably, while tissue-resident AMs self-renew at steady state, following infections or injury resident AMs are replaced by monocyte-derived AMs that differ from resident AMs phenotypically and metabolically. Immature AMs have been reported in several infections and lung pathologies, including fibrosis, but not yet in lung cancer. Here we demonstrate that during lung cancer the residential AM pool is replaced by monocyte-derived AMs exhibiting a profibrotic and immature-like state. Moreover, these cells exhibit an inflammatory signature distinct from residential AMs. In a lung cancer setting, Siah1a and Siah2 loss enhances the immature state of monocyte-derived AMs and confers on them a unique profibrotic signature favoring lung cancer progression.

We also show that genes significantly upregulated in Siah1a/2-deleted immature-like AMs are associated with a poor survival in patient with lung adenocarcinoma. Notably, AI-based deconvolution of scRNAseq analyses of LUAD patients carrying the gene signature identified in the Siah1a/2 ablated immature-like AMs revealed an association with higher infiltration of CD14^+^ myeloid cells and worse survival. This important observation implicates the monocytes/ macrophages infiltrated cell population in lung cancer progression. Among the genes upregulated in Siah1a/2-deleted immature-like AMs and associated with the highest significant hazard score were those involved in profibrotic macrophages. Consistent with our findings are reports implicating TAMs as important contributors to metabolism and matrix processing, which underlie lung cancer progression ^46, 47, 48, 49^. Along these lines, pulmonary fibrosis is a risk factor for developing lung carcinogenesis, and patients with idiopathic pulmonary fibrosis (IPF) later diagnosed with lung cancer tend to have a poorer prognosis than patients with lung cancer without IPF, regardless of treatment modality.

Overall, these studies indicate that Siah1a/2 regulates AM differentiation and activity in a manner that controls immune cell recruitment and tumor growth. These findings could suggest new approaches to monitor or target specific macrophage populations, as means to antagonize lung cancer. Understanding molecular mechanisms underlying AM differentiation and control of lung cancer initiation and progression could offer better monitoring, stratifications and treatment options for patients with lung cancer.

## Acknowledgements

We thank members of the Ronai lab for discussion and Shared Resources for various analyses (vivarium, FACS, bioinformatics) supported by P30CA030199 grant. Support by R35CA197465 to ZR is gratefully acknowledged.

## Author contributions

MS and ZR designed the study. MS and YD performed the experiments. LB helped with immune study design and evaluation. RM, ER, KW performed bioinformatic analyses. MS and ZAR wrote the manuscript.

## Conflict of Interest

ZAR is co-founder and scientist advisor of Pangea Biomed. No conflict of interest is reported for any of the other authors.

## Material and Methods

### Animals and tumor model

All experimental animal procedures were approved by the Institutional Animal Care and Use Committee of Sanford Burnham Prebys Medical Discovery Institute. *Siah1a*^*f/f*^ *and Siah2*^*f/f*^ were generated by Cyagen. *Lyz2*^*Cre*^ and *CD11c*^*Cre*^ mice were purchased by Jackson. For the urethane-induced lung cancer model, mice were injected intraperitoneally with urethane (Sigma, 1mg/gr mouse) once a week for 6 weeks. Lungs were collected either 7 or 24 weeks after first urethane injection.

### Bone marrow chimeras

C57Bl6 recipient mice were lethally-irradiated (1000 rads) and reconstituted by intravenous (i.v.) injection of 1 × 10^7^ bone marrow (BM) cells isolated from femurs and tibias of donor *WT CD45*.*2+, cSiah1a/2*^*f/f*^*::Lyz2*^*Cre*^ *CD45*.*2+* or *WT CD45*.*1+* mice. Irradiated mice were injected intravenously with a 1:1 mixture of either *WT CD45*.*2+ and WT CD45*.*1+* cells or *cSiah1a/2*^*f/f*^*::Lyz2*^*Cre*^ *CD45*.*2*^*+*^ cells and *WT CD45*.*1*^*+*^ cells for a total 1 × 10^7^ BM cells. Recipients were treated with antibiotics (trimethoprim 8 mg/ml and sulfamethoxazole 40 mg/ml in drinking water) for 3 weeks after injection. Reconstitution was confirmed 6–8 weeks after BM transfer.

### Tissue digestion

Tissues were excised, minced, and digested with 1 mg/ml collagenase D (Roche) and 100 µg/ml DNase I (Sigma) at 37°C for 45 minutes. Digests were then passed through a 70-μm cell strainer to generate a single-cell suspension. Cells were washed twice with PBS containing 2 mM EDTA and stained for flow cytometry.

### Flow cytometry

Tissue-derived single-cell suspensions were washed twice with FACS staining buffer, fixed 15 min with 1% formaldehyde in PBS, washed twice, and resuspended in FACS staining buffer for staining with specific antibodies, followed by fixation with 1% formaldehyde and then flow cytometry analysis. The following antibodies were purchased from Biolegend: CD45.2 (104), CD8a (53-6.7), CD4 (GK1.5), CD45.1 (A20), CD11c (N418), CD11b (M1/70), MHC class I (AF6-88.5), CD80 (16-10A1), GR1 (RB6-8C5), CD206 (C068C2), F4/80 (BM8), SiglecF (S17007L), Ly6C (HK1.4), Ly6G (1A8), NK1.1 (PK136), B220 (RA3-6B2), CD44 (IM7). All data were collected on an LSRFortessa cell analyzer (BD Biosciences) and analyzed using FlowJo Software (Tree Star).

### Histology and immunofluorescence

Lungs were fixed in 4% formalin overnight at 4°C, washed with PBS, paraffin-embedded, cut into 5 μm-thick sections and stained with H&E. For immunofluorescence, sections were deparaffinized, rehydrated and washed in PBS. Antigen retrieval was performed in a pressure cooker (Decloaking chamber, Biocare Medical) in citrate buffer (pH 6.0). Ki67 (AbCam Ab15580) immunostaining was performed by incubating sections overnight at 4°C with antibodies in Dako antibody diluent. Alexa Fluor 594-conjugated secondary antibodies was added for 1h at room temperature (Molecular Probes), and nuclei were counterstained using SlowFade Gold Antifade reagent (Vector) with 4′,6-diamidino-2-phenylindole (DAPI, Vector). Image data were obtained using Olympus TH4– 100 microscope and using Slidebook 4.1 digital microscopy. For quantification Ki67, positive cells were counted in ten random × 20 fields per mouse. Staining was scored by counting Ki67-positive relative to total cells.

### Immunofluorescence

To stain AMs grown in culture, cells were grown on coverslips and fixed in fixation buffer (4% paraformaldehyde/2% Sucrose/PBS) for 20 min at room temperature. Coverslips were then rinsed twice in PBS and permeabilized in permeabilization buffer (0.4% Triton-X and 1% BSA in PBS) for 20 min. Primary antibodies were applied at 1:250 dilution in staining buffer (0.1% Triton-X and 0.1% BSA in PBS) overnight at 4°C in a humid chamber. Coverslips were washed 5 times (5-min each) in wash buffer (0.2% Triton-X and 0.2% BSA in PBS). Secondary antibodies (AlexaFluor secondary 488 or 594, Invitrogen) were applied at 1:250 dilution in staining buffer for 2-3 hours at room temperature in a humid chamber in the dark. Prior to mounting with Vectorshield with DAPI (Vector Laboratories, CA), coverslips were washed twice more in wash buffer. Immunofluorescent analysis was conducted on an Olympus TH4-100 fluorescent microscope using Slidebook V.4.1 digital microscopy.

### Quantification of lung cancer

Fixed and embedded lung tissues were sliced to obtain five serial sections per lung. H&E staining was used to quantify number and size of nodules under a histology microscope. Size was calculated as the area of nodules using a measurement tool provided by CellSens Standard v3.2 software (Olimpus America).

### AM primary cell culture

Lung tissue was digested as described above to obtain a single cell suspension. SiglecF-positive cells were isolated using anti-SiglecF microbeads (Miltenyi Biotec) and plated on non-treated 6-well plates in RPMI 1640 media containing 1x Glutamax, 1x Pyruvate, 1x penicillin/Streptomycin, 10% FBS and 20 ng/ml of recombinant GM-CSF, based on a previous publication ^50^. After 16 hours media was replaced with fresh culture media, which was replaced with new media every 2 days. Cells were collected either the day after plating or after 6 days. When indicated, AMs were treated with 3uM XAV939 (Sigma) or 3uM ML385 (MedChem Express) overnight.

### Bone marrow-derived cell culture

Mice were euthanized with CO_2_, the femurs were removed, and BM cells were harvested and washed. For macrophage differentiation, BM cells were resuspended in RPMI-1640 medium containing 10% FBS, penicillin/streptomycin, and 2 mM L-glutamine and placed in Petri dishes. L929 cell supernatants (a source of macrophage-colony stimulating factor, M-CSF) were added at 30% (vol/vol), and cells were incubated 7 days. Differentiated macrophages were then harvested and used in the experiments.

### Western blotting

Cells were washed once with PBS at room temperature and resuspended in RIPA buffer (PBS containing 1% NP-40, 1% sodium deoxycholate, 1% SDS, 1 mM EDTA, and phosphatase and protease inhibitors), while tissue was homogenized directly in RIPA buffer. Lysates were centrifuged, and supernatants were removed and subjected to SDS-PAGE. Proteins were transferred to nitrocellulose membranes (Osmonics Inc., MN, USA), which were blocked and incubated with respective primary antibodies followed by Alexa Fluor-conjugated or HRP-conjugated secondary antibodies. Blots were imaged using an Odyssey detection system (Amersham Bioscience, NJ, USA) or a ChemiDoc MP imaging system (BioRad).

### RNA extraction and qRT-PCR analyses

Total RNA was extracted from tumors or cells using TRIzol (Ambion) and treated with DNase I. cDNA was synthesized using oligo-dT and random hexamer primers according to the SYBR Green qPCR protocol (Life Technologies). Total RNA was reverse transcribed using High Capacity Reverse Transcriptase kits (Invitrogen), according the manufacturer’s protocol. RNA purity and concentration was checked and quantified by reading absorbance at 260 and 280 nm in a NanoDrop spectrophotometer (Thermo Fisher). qRT-PCR analyses were performed using SYBR Green RT-PCR kits (Invitrogen) on a Bio-Rad CFX Connect Real-Time system or Roche LightCycler. GAPDH or 18S served as internal control. PCR primers were designed using Primer3, and their specificity was checked using BLAST. PCR products were limited to 100–200 bp. Primer sequences are shown in Supplemental Table 2.

### RNA-seq analysis

Lung tissues from mutant or WT mice were digested as described above, and CD11c+ SiglecF+ F4/80+ cells were sorted by flow cytometry and processed for gene expression analysis. PolyA RNA was isolated using the NEBNext® Poly(A) mRNA Magnetic Isolation Module, and bar-coded libraries were constructed using the NEBNext® Ultra™ Directional RNA Library Prep Kit for Illumina® (NEB, Ipswich MA). Libraries were pooled and sequenced single-end (1×75) on an Illumina NextSeq 500 using the High output V2 kit (Illumina Inc., San Diego CA). The RNA-seq samples were sequenced at a sequencing depth of 11-16 million reads.

Illumina Truseq adapters and polyA/polyT sequences were trimmed from raw reads using Cutadapt version 2.3 ^51^, Trimmed reads were aligned to mouse genome version mm10 and Ensembl gene annotations version 84 using STAR version 2.7.0d_0221 ^52^ adopting alignment parameters from ENCODE long RNA-seq pipeline (https://github.com/ENCODE-DCC/long-rna-seq-pipeline). RSEM version 1.3.1 ^53^ was used to obtain gene level estimated counts and transcripts per million (TPM). FastQC version 0.11.5 (https://www.bioinformatics.babraham.ac.uk/projects/fastqc/) and MultiQC version 1.8 ^54^ were used to assess the quality of trimmed raw reads and alignment to genome as well as transcriptome. To remove lowly expressed genes from downstream analysis, only genes with RSEM estimated counts equal or greater than 5 times the total number of samples were retained for differential expression analysis. Differential expression comparisons were performed using Wald test implemented in DESeq2 version 1.22.2 ^55^. Genes with Benjamini-Hochberg corrected p-value < 0.05 and fold change ≥ 1.5 or ≤ -1.5 were identified as differentially expressed. Pathway analysis was performed in Ingenuity Pathway Analysis (Qiagen, Redwood City, USA).

### Single-cell library preparation and sequencing

Mutant or WT mice were sacrificed 7 weeks after the first urethane treatment. Lungs were digested as described above. Single cell suspensions were washed with 1XPBS-4%FBS before incubation for 20 min on ice at 5•10^7^ cells/ml with 500 ng/ml Fc block (2.4G2, BD Pharmingen). Cells were then incubated for 1 hour on ice with AF700 conjugated CD45.2 monoclonal antibody (104, Biolegend). For scRNA-seq libraries, DAPI negative (live) CD45+ cells were sorted using a flow cytometer, and sorted cells were resuspended in RPMI for counting. Libraries were prepared using Single Cell 3’ Reagent Kits v3.1: Chromium™ Single Cell 3’ Library & Gel Bead Kit v3.1, PN-1000196 and PN-1000129 and following the Single Cell 3’ Reagent Kits v3 (PN-1000269) User Guide (Manual Part # CG000315 Rev C). Libraries were sequenced on NovaSeq S4, using half full lanes for 2 samples, with mean read depth of 87-128 thousand reads per cell.

### scRNA-seq data pre-processing

10X Genomics Cell Ranger pipeline version 6.1.2 was used for processing the scRNA-seq datasets. Single cell gene counts for each sample were generated using *cellranger count* and mouse genome version mm10 annotated with mouse GENCODE version M28 ^56^. There were 8,522 cells obtained for *Siah* WT (128 thousand mean reads per cell, 2674 median genes detected per cell) and 6,479 cells for *Siah2*^*-/-*^ (103 thousand mean reads per cell, 2502 median genes detected per cell). The gene count matrices for *Siah* WT and *Siah2*^*-/-*^ were aggregated and normalized for effective sequencing depth using *cellranger aggr* applying “--normalize=mapped” parameter.

### Integrated analysis of scRNA-seq datasets

Aggregated gene count matrix for *Siah* WT and *Siah2*^*-/-*^ scRNA-seq samples were processed using Seurat version 4.0.5 ^57^ and R version 4.0.2. 10X Genomics Cell Ranger aggr matrix was converted to Seurat object by retaining genes expressed in minimum of 10 cells and cells expressing a minimum of 200 genes. Cells expressing <10% mitochondrial genes (to remove dead/low quality cells) and <5000 genes expressed (to remove potential doublets/multiplets) were retained. Integration of *Siah* WT and *Siah2*^*-/-*^ samples was performed using *sctransform* normalization method (https://satijalab.org/seurat/articles/integration_introduction.html#performing-integration-on-datasets-normalized-with-sctransform-1). 3000 variable genes were used in *SelectIntegrationFeatures* step. The two datasets were integrated using *FindIntegrationAnchors(normalization*.*method = “SCT”)* and *IntegrateData(normalization*.*method = “SCT”)*. PCA components were computed using *RunPCA()*. Cell Clusters were computed using *RunUMAP(dims=1:30), FindNeighbors()*, and *FindClusters(resolution=0*.*5)* resulting in 17 cell clusters. For visualization of gene expression and differential expression analysis, default assay was set to *“RNA”* and gene counts normalized using *NormalizaData()*. Cluster markers were found using *FindAllMarkers()*. Differential expression analysis comparisons were performed using *FindMarkers(test*.*use = “MAST”)*. Plots were prepared using Seurat and ggplot2. Pathway analyses were performed in Ingenuity Pathway Analysis (Qiagen, Redwood City, USA). RNA-seq and scRNA-seq main and supplemental plots were prepared using ggplot2 (H. Wickham. ggplot2: Elegant Graphics for Data Analysis. Springer-Verlag New York, 2016), ComplexHeatmap ^58^, and Seurat.

### LUAD analysis

Gene expression (RNA-seq) and clinical data from TCGA-LUAD Pan-Cancer 2018 ^59^ were downloaded from cBioPortal ^60^. Survival analysis was performed in R version 4.2.1 using survival (Therneau T (2020). A Package for Survival Analysis in R. R package version 3.3-1, URL: https://CRAN.R-project.org/package=survival.), survminer (Alboukadel Kassambara, Marcin Kosinski and Przemyslaw Biecek (2020). survminer: Drawing Survival Curves using ‘ggplot2’. R package version 0.4.9. http://www.sthda.com/22nglish/rpkgs/survminer/), and maxstat (Torsten Hothorn (2017). Maxstat: Maximally Selected Rank Statistics. R package version 0.7-25. https://CRAN.R-project.org/package=maxstat) packages. Categorization of TCGA-LUAD samples as ‘high’ and ‘low’ gene expressors in TCGA data was determined using surv_cutpoint() and surv_categorize() functions from survminer package. Survival analyses using gene signatures (multiple genes) were performed by computing mean expression of the gene signature. Hazard ratios in TCGA-LUAD for each gene were computed in R with coxph() function using the same ‘high’ and ‘low’ gene expression categorizations described above. Wang et al. developed CODEFACS (COnfident DEconvolution For All Cell Subsets), a tool deconvolving cell-type-specific gene expression in each sample from bulk expression given either precomputed estimates of cell abundance or cell-type-specific signature^44^. To deconvolve TCGA-LUAD and TCGA-SCC, they first estimated abundance of 11 cell types (macrophages/dendritic cells--CD14+, B cells--CD19+, CD4+T cells, CD8+ T cells, T regulatory cells, NK cells--CD56+, endothelial cells, fibroblasts, neutrophils, basophils, eosinophils and tissue-specific tumor cells) based on bulk methylation and then applied CODEFACS to the corresponding bulk gene expression. The output is cell-type-specific gene expression profile for each sample in TCGA-LUAD and TCGA-SCC.

### Statistical Analysis

GraphPad Prism version 7 was used for statistical analysis. Differences between two groups were assessed using two-tailed unpaired *t* test. Differentially expressed genes were identified in bulk RNA-seq using Wald test implemented in DESeq2. Differentially expressed genes in scRNA-seq clusters were identified using *FindMarkers()* and *MAST* test in Seurat. Survival analysis and hazard ratio p-values were computed using log-rank test. Differences in cell type compositions of TCGA-LUAD patients were tested using Wilcoxon rank sum test.

## Supplemental Figures

**Sup. Figure 1.**
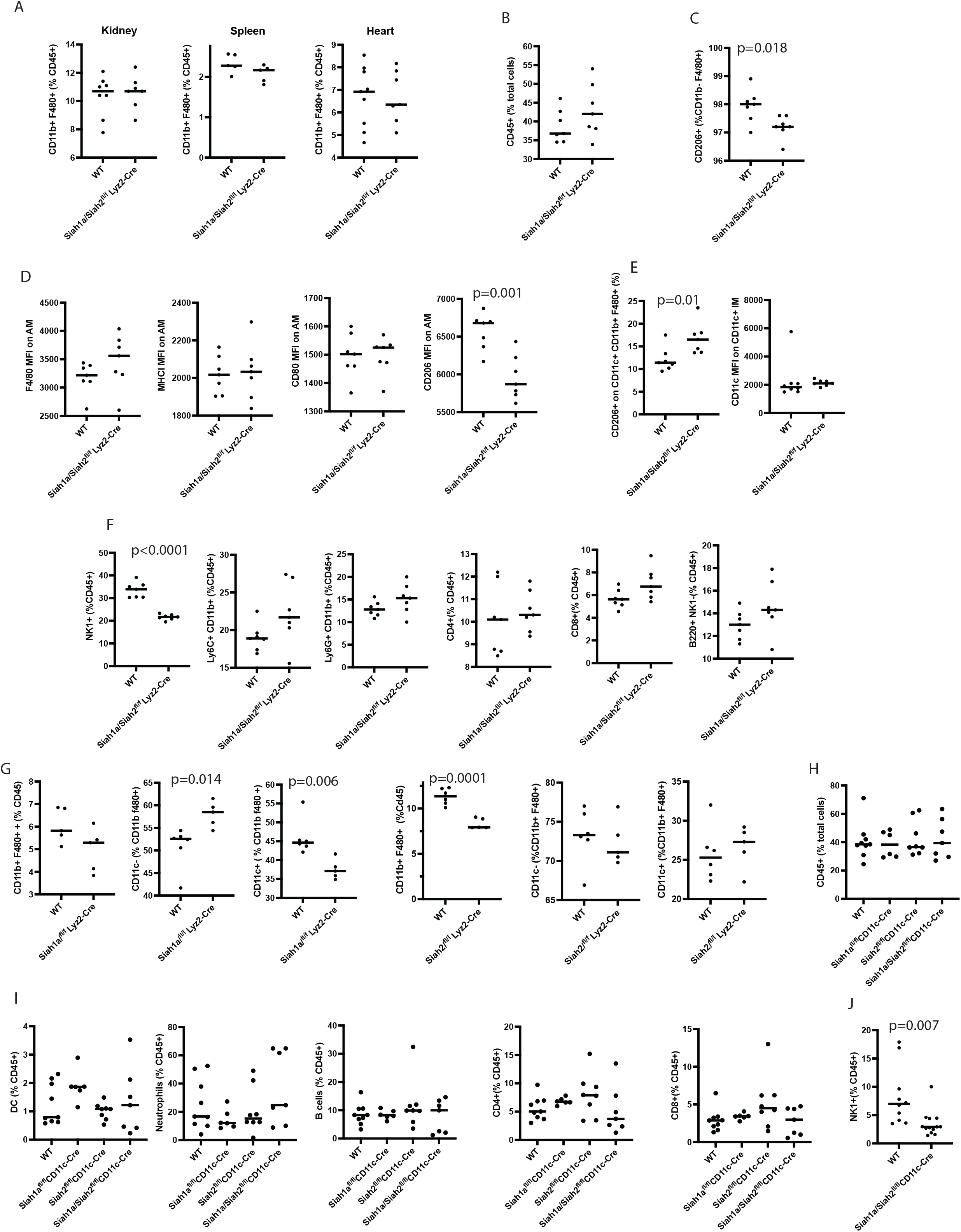
Siah1a/2-deleted macrophages exhibit a specific immune phenotype in lung tissue. **A** Frequency of F480^+^ CD11b^+^ immune cells among CD45^+^ cells from kidney, spleen and heart. n=8 for WT; n=7 for *Siah1a*^*f/f*^ : *Siah2*^*f/f*^ : *Lyz2-Cre*. **B** Quantification of CD45+ cells from lungs of *WT* and *cSiah1a/2*^*f/f*^*::Lyz2*^*Cre*^ mice at a homeostatic state. Quantification is shown as a percentage of immune cells among total cells. n=7 for WT; n=7 for *cSiah1a/2*^*f/f*^*::Lyz2*^*Cre*^. **C** Frequency of CD206^+^ cells among F480^+^ CD11b low cells from lung. n=7 for both genotypes. **D** Expression of F4/80, MHCI, CD80, and CD206 on AMs (CD11b low F480^+^ CD11c^+^ SiglecF^+^ cells) based on flow cytometry analysis of lungs from *WT* and *cSiah1a/2*^*f/f*^*::Lyz2*^*Cre*^ mice. n=7 for WT; n=7 for *cSiah1a/2*^*f/f*^*::Lyz2*^*Cre*^. **E** Frequency of CD206+ cells and expression of CD11c among CD11c^+^ IMs of lungs of WT and *cSiah1a/2*^*f/f*^*::Lyz2*^*Cre*^ mice. n=7 for both genotypes. **F** Frequency of NK1^+^, LY6C^+^ CD11b^+^, LY6G^+^ CD11b^+^, CD4^+^, CD8^+^, and B220^+^ NK1^-^ immune cells among CD45^+^ cells from lungs of WT and *cSiah1a/2*^*f/f*^*::Lyz2*^*Cre*^ mice. n=7 for both genotypes. **G** Frequency of CD11b^+^ and F4/80^+^ cells among CD45^+^ cells, and frequency of CD11c^+^ and CD11c^-^ among IMs of lungs of WT and *cSiah1a*^*f/f*^*::Lyz2*^*Cre*^ mice or *cSiah2*^*f/f*^*::Lyz2*^*Cre*^ mice. n=6 for WT, n = 5 for *cSiah1a*^*f/f*^*::Lyz2*^*Cre*^; n = 6 for WT, n = 6 for *cSiah2*^*f/f*^*::Lyz2*^*Cre*^. **H** Quantification of CD45+ cells of *WT, cSiah1a*^*f/f*^*::CD11c*^*Cre*^, *cSiah2*^*f/f*^*::CD11c*^*Cre*^ and *cSiah1a/2*^*f/f*^*::CD11c*^*Cre*^ mice. Quantification is shown as a percentage of indicated immune cells among total cells. n=7 for WT; n=5 for *cSiah1a*^*f/f*^*::CD11c*^*Cre*^; n=8 for *cSiah2*^*f/f*^*::CD11c*^*Cre*^; n=6 for *cSiah1a/2*^*f/f*^*::CD11c*^*Cre*^. **I** Quantification of dendritic cells (DCs), neutrophils, B cells, and CD4+ or CD8+ cells from *WT, cSiah1a*^*f/f*^*::CD11c2*^*Cre*^, *cSiah2*^*f/f*^*::CD11c*^*Cre*^ and *cSiah1a/2*^*f/f*^*::CD11c*^*Cre*^ mice. Quantification is shown as a percentage of indicated immune cells among CD45.2+ cells. n=7 for WT; n=5 for *cSiah1a*^*f/f*^*::CD11c*^*Cre*^; n=8 for *cSiah2*^*f/f*^*::CD11c*^*Cre*^ n=6 for *cSiah1a/2*^*f/f*^*::CD11c*^*Cre*^. **J** Quantification of NK1+ cells from *WT* and *cSiah1a/2*^*f/f*^*::CD11c*^*Cre*^ mice. Quantification is shown as a percentage of indicated immune cells among CD45.2^+^ cells. n=11 for WT; n=13 for *cSiah1a/2*^*f/f*^*::CD11c*^*Cre*^. Data were analyzed by unpaired t-test.

**Sup. Figure 2.**
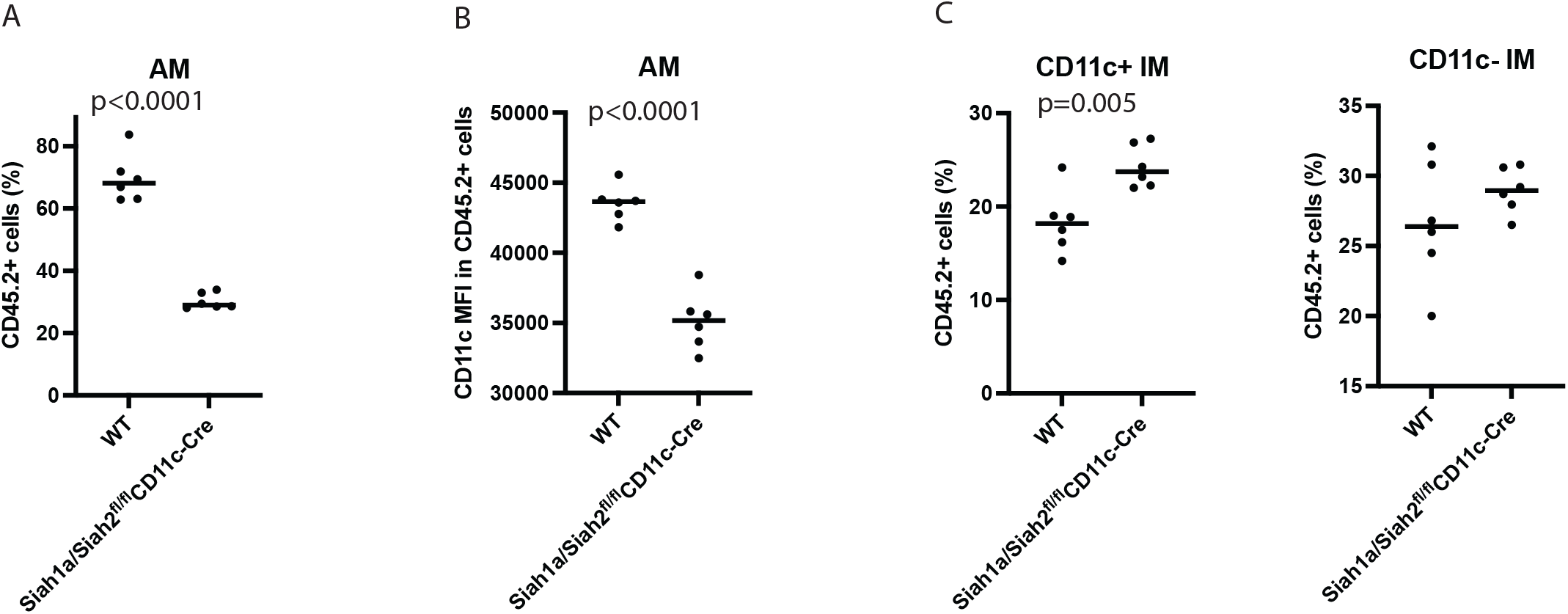
Frequency of selected immune cells in lungs of transplanted mice. Bone marrow cells from *cSiah1a/2*^*f/f*^*::CD11c*^*Cre*^ (CD45.2^+^) or CD45.2 WT mice were co-transferred with CD45.1 competitors bone marrow cells into lethally-irradiated C57Bl6 mice, and lungs were harvested 10 weeks later. **A** Flow cytometry quantitation of CD45.2 positive cells on lung AMs (CD11b low F4/80^+^ CD11c^+^ SiglecF^+^) from irradiated mice co-transfected with either a mix (1:1) of *cSiah1a/2*^*f/f*^*::CD11c*^*Cre*^ CD45.2^+^ : CD45.1^+^ cells (n=6) or CD45.2 WT: CD45.1^+^ cells (n=6). **B** CD11c expression on CD45.2^+^ AMs of *cSiah1a/2*^*f/f*^*::CD11c*^*Cre*^ CD45.2^+^ : CD45.1^+^ or CD45.2 WT: CD45.1^+^ co-transfected mice. n=6 for both genotypes. **C** Flow cytometry quantitation of CD45.2^-^ positive cells on lung CD11c^+^ and CD11c^-^ IMs (CD11b^+^ F4/80^+^ CD11c^-^) from irradiated mice co-transfected with either a mix (1:1) of *cSiah1a/2*^*f/f*^*::CD11c*^*Cre*^ CD45.2^+^ : CD45.1^+^ cells (n=6) or CD45.2 WT: CD45.1^+^ cells (n=6). Data were analyzed by unpaired t-test.

**Sup. Figure 3.**
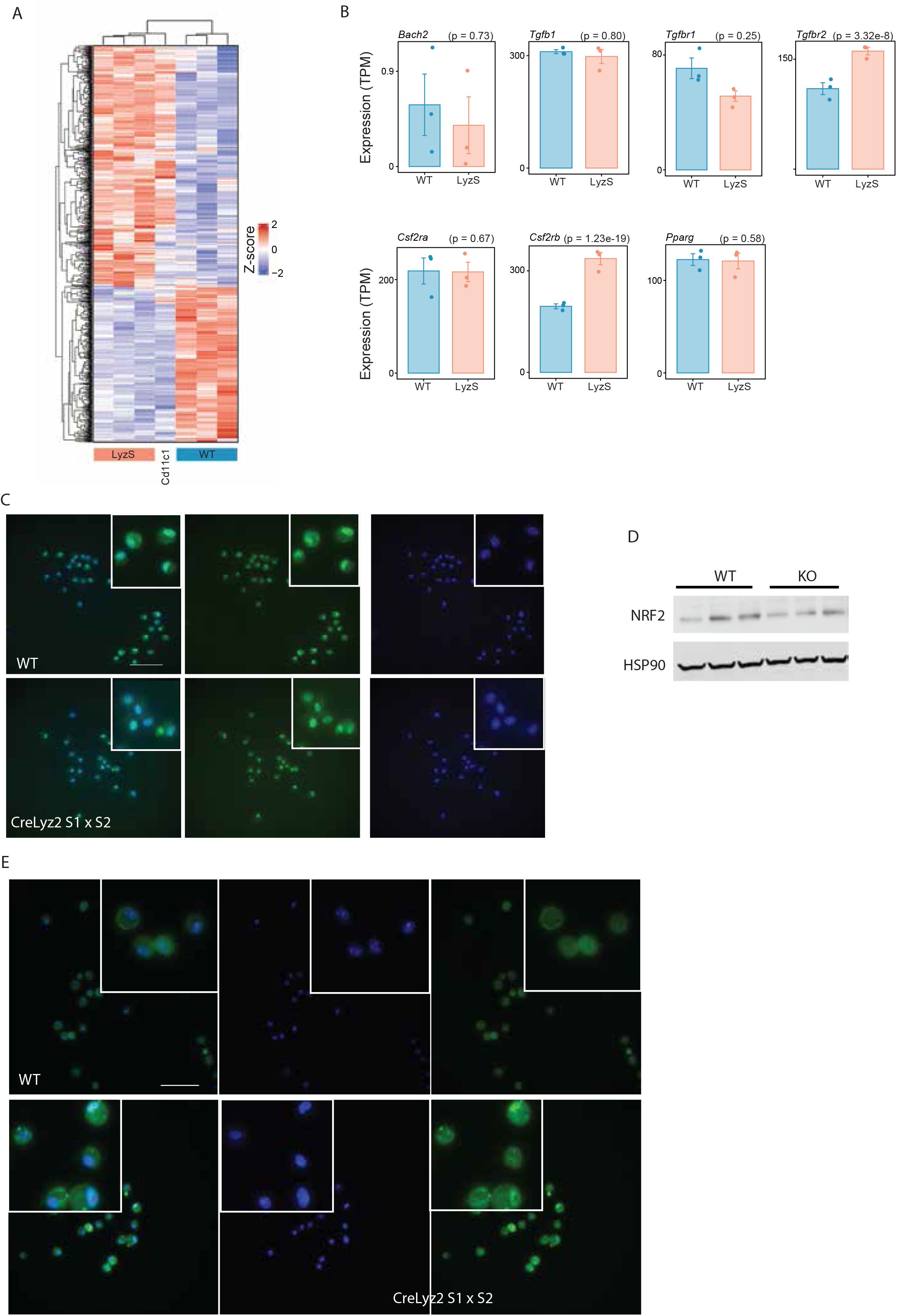
Siah1a/2 deletion in AMs promotes β catenin nuclear localization. RNA-seq analysis of CD11c^+^ F480^+^ SiglecF^+^ cells sorted from lungs of WT (n=3), *cSiah1a/2*^*f/f*^*::Lyz2*^*Cre*^ (n=3), *cSiah1a/2*^*f/f*^*::CD11c*^*Cre*^ (n=1) mice. Each sample was a pool of 3 lungs. **A** Heatmap showing differentially expressed genes in *cSiah1a/2*^*f/f*^*::Lyz2*^*Cre*^ vs. WT comparison. Also shown is expression fo the same genes in *cSiah1a/2*^*f/f*^*::CD11c*^*Cre*^ indicating that *cSiah1a/2*^*f/f*^*::Lyz2*^*Cre*^ and *cSiah1a/2*^*f/f*^*::CD11c*^*Cre*^ AMs show similar gene expression patterns. **B** Expression levels of selected genes from WT and *Siah1a*^*f/f*^ : *Siah2*^*f/f*^ : *Lyz2-Cre* AMs based on RNA-seq gene expression data. n = 3 for both genotypes. **C** PPARγ staining of primary AMs cultured overnight in media containing GM-CSF. PPARγ = green, Dapi = blue. n =3 for each genotype. Scale bar, 600 µM. **D** Primary bone marrow-derived macrophages (BMDMs) were cultured 9 days for western blot analysis. Lysates from WT and *cSiah1a/2*^*f/f*^*::Lyz2*^*Cre*^ BMDMs were immunoblotted for NRF2. HSP90 was used as a loading control. n = 3 for both genotypes. **E** Staining for non-phosphorylated Ser45 (active) β−catenin in AMs cultured overnight in media containing GM-CSF and quantification. β-catenin = green, Dapi = blue. n =3 for each genotype. Scale bar, 600 µM. Data in panel B were analyzed by Wade test.

**Sup. Figure 4.**
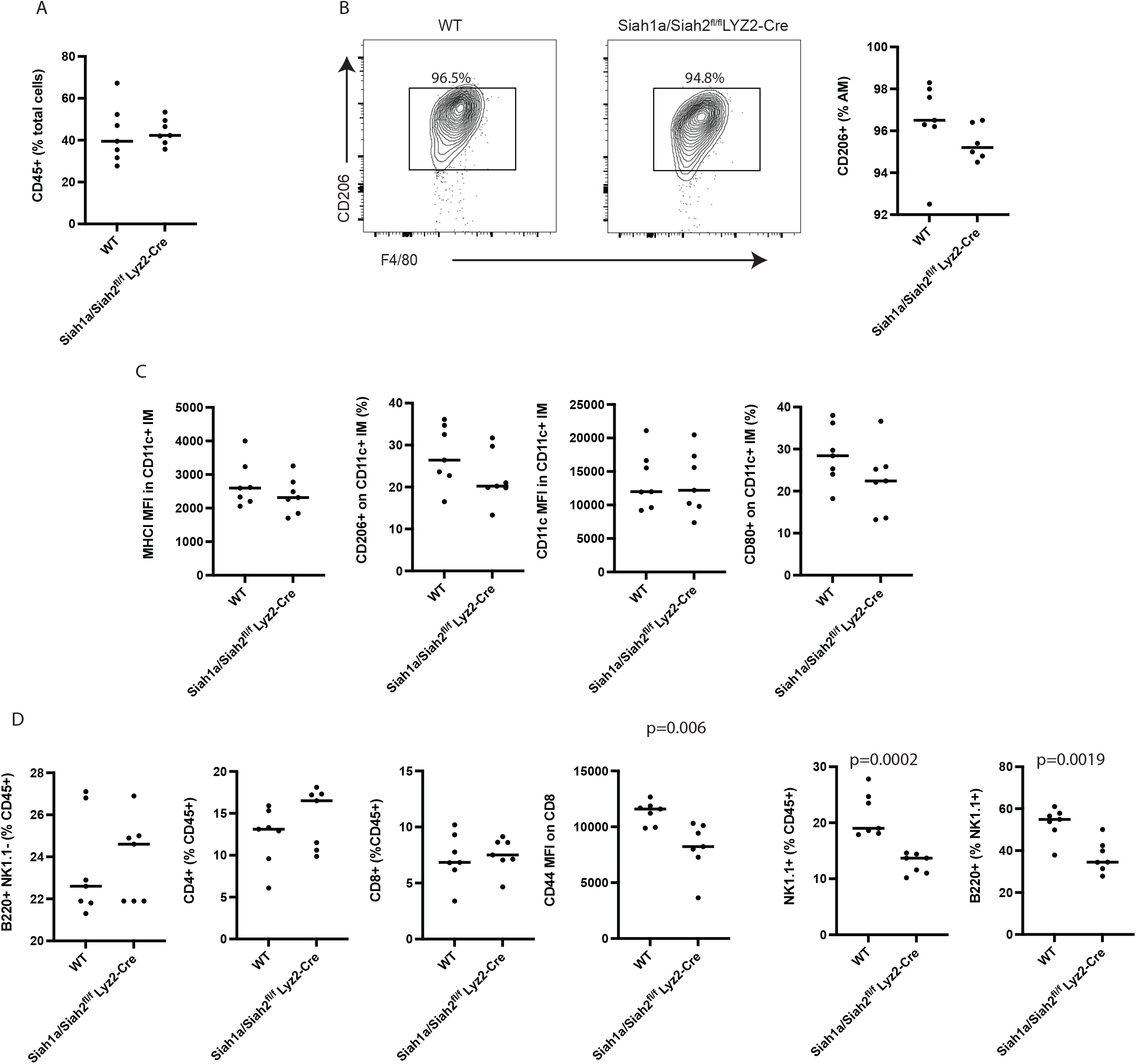
Siah1a/2 deletion in macrophages promotes lung cancer following urethane treatment. Urethane (1 mg/g of mouse weight) was injected intraperitoneally once a week for 6 weeks, and lungs were collected 22 weeks later. **A** Frequencies of CD45^+^ cells among total cells of lungs after urethane injection. n = 7 for each genotype. **B** Representation and quantitation of CD206^+^ cells among AMs. Quantification is shown as a percentage of indicated immune cells among AMs^+^ cells. n = 7 for each genotype. **C** Expression of MHCI, CD206 and CD11c and frequency of CD80 in the CD11c^+^ IM population. n = 7 for each genotype. **D** Frequencies of B (B220^+^ NK1^-^), CD4^+^, CD8^+^ NK1.1^+^ cells among CD45^+^ cells, expression of CD44 on CD8^+^ cells and frequency of activated NK1.1+ cells (B220^+^) among NK1.1^+^ cells. n = 7 for each genotype. Data were analyzed by unpaired t-test.

**Sup. Figure 5.**
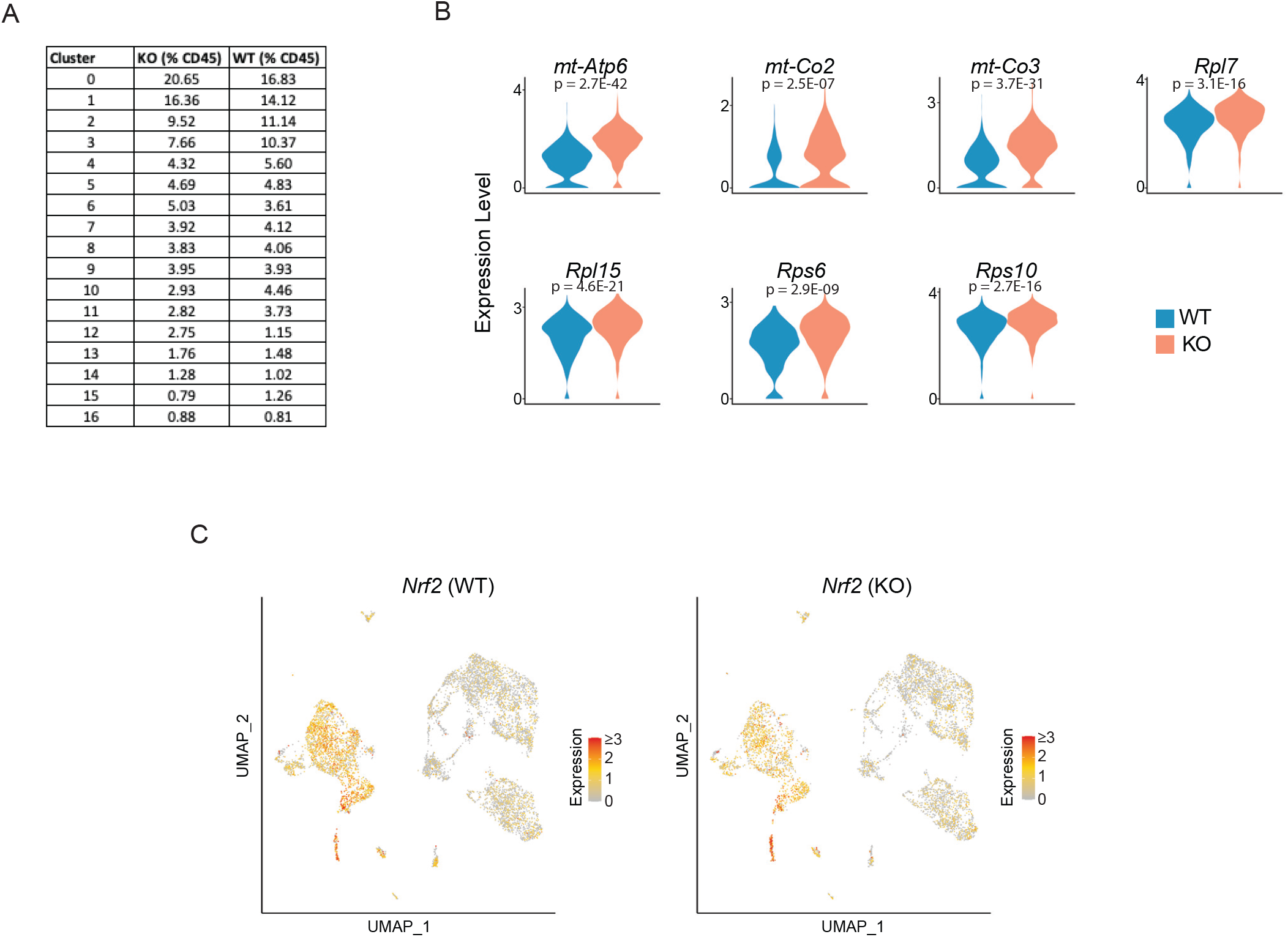
Single cell RNA-seq analysis of CD45+ cells in lungs of urethane-treated mice. CD45^+^ cells were sorted by flow cytometry from WT and *cSiah1a/2*^*f/f*^*::Lyz2*^*Cre*^ lungs collected 7 weeks after the first urethane injection, and single cell RNA-seq was performed. **A** Table showing the proportion of cells in each cluster within CD45^+^ clusters from WT and *cSiah1a/2*^*f/f*^*::Lyz2*^*Cre*^ CD45^+^ cells, related to UMAP in Figure 6A. **B** Violin plots comparing expression levels of select genes from cluster 5 (macrophages). **C** UMAP plots showing expression of NRF2 in all CD45+ cell clusters. Data in panel B were analyzed using *FindMarkers()* and *MAST* test in Seurat.

**Sup. Figure 6.**
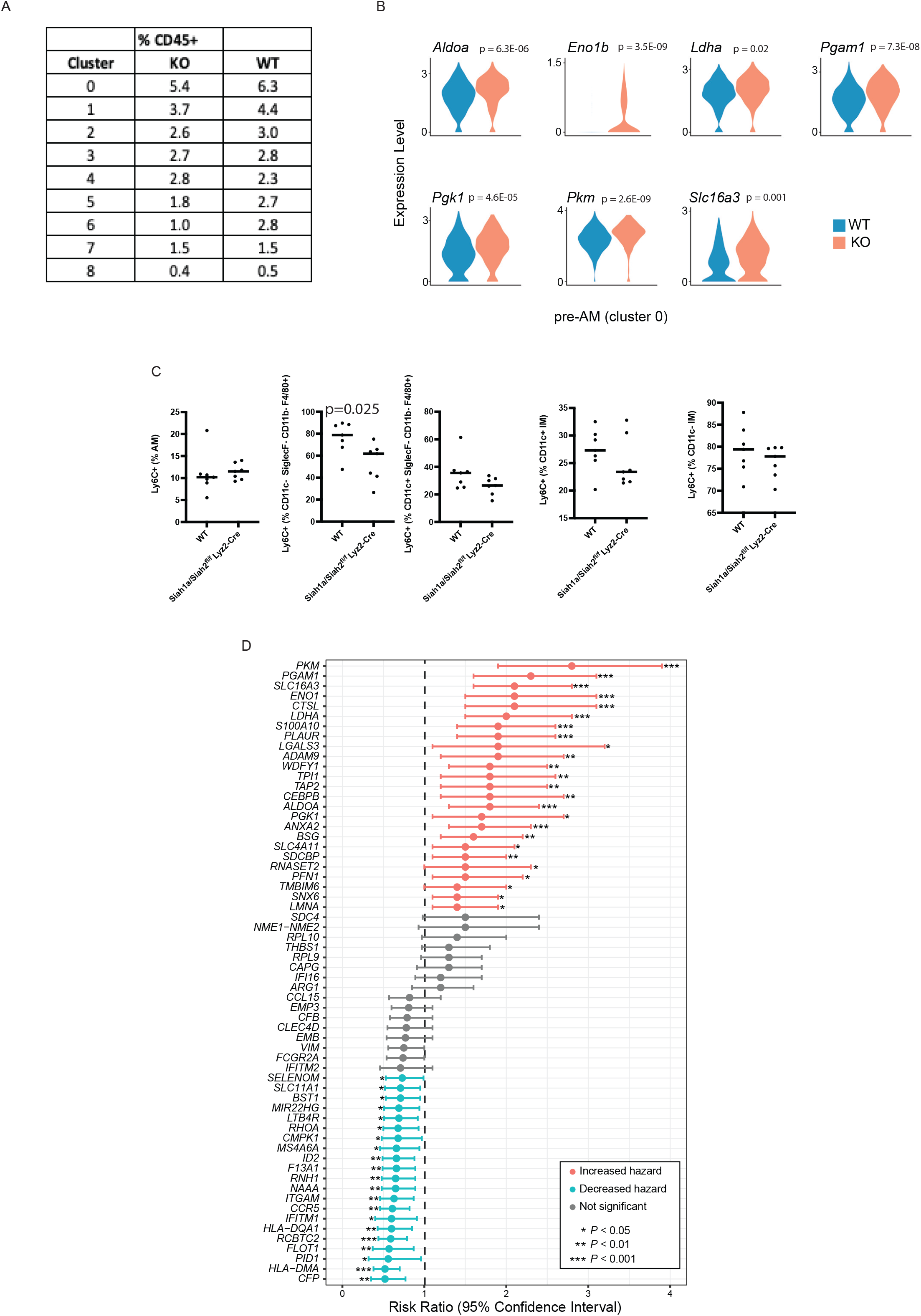
Single cell RNA-seq analysis of myeloid cells in lungs of urethane-treated mice. Clusters 2, 3 and 5 from single cell RNA-seq analysis of CD45+ cells (Figure 6A) containing myeloid cells were re-clustered to provide higher resolution clustering into 8 new clusters. **A** Table representing the proportion of cells within each myeloid cell cluster from WT and *cSiah1a/2*^*f/f*^*::Lyz2*^*Cre*^ cells. **B** Violin plots comparing expression of select genes from sub-cluster 0 in myeloid cells. **C** Frequencies of LY6C^+^ cells among mature AMs, immature AMs (CD11c^-^ SiglecF^-^ CD11b low and F480^+^ and CD11c^+^ SiglecF^-^ CD11b low and F480^+^), CD11c^+^ IMs and CD11c^-^ IMs in lungs at 22 weeks after urethane injection. n = 6 for each genotype. **D** 72 Inflammatory genes significantly upregulated in myeloid sub-cluster 0 of Siah1a/2-deleted cells relative to WT were used to calculate hazard ratios in the TCGA-LUAD patient cohort. For each gene, the cohort was stratified as high and low expressors using optimally best cutoff. Shown are hazard ratios, 95% confidence intervals, and p-values (log-rank test). Data in panel B were analyzed using *FindMarkers()* and *MAST* test in Seurat, data in panel C were analyzed by unpaired t-test.

## Notes

### Competing Interest Statement

ZAR is co-founder and scientific advisor of Pangea BioMed

